# Metabolic tuning enables immediate adaptation to energy stress in yeast

**DOI:** 10.1101/2025.06.06.658098

**Authors:** Katherine Bexley, Michaela Ristová, Sushma Sharma, Christos Spanos, Andrei Chabes, David Tollervey

**Affiliations:** Institute of Cell Biology, University of Edinburgh, Edinburgh, Scotland; Dept. of Medical Biochemistry and Biophysics, Umeå University, Umeå Sweden

**Keywords:** stress, yeast, gene expression, translation regulation, metabolomics, RNA-protein interaction

## Abstract

In *Saccharomyces cerevisiae*, glucose depletion induces metabolic reprogramming through widespread transcriptional and translational reorganization. We report that initial, very rapid translational silencing is driven by a specialized metabolic mechanism. Following glucose withdrawal, intracellular NTP levels drop drastically over 30 sec, before stabilizing at a regulated, post-stress set-point. Programmed translational control results from the differential NTP affinities of key enzymes; ATP falls below the (high) binding constants for DEAD-box helicase initiation factors, including eIF4A, driving mRNA release and blocking 80S assembly. Contrastingly, GTP levels always greatly exceed the (low) binding constants for elongation factors, allowing ribosome run-off and orderly translation shutdown. Translation initiation is immediately lost on all pre-existing mRNAs, before being preferentially re-established on newly synthesized, upregulated stress-response transcripts. We conclude that enzymatic constants are tuned for metabolic remodeling. This response counters energy depletion, rather than being glucose-specific, allowing hierarchical inhibition of energy-consuming processes on very rapid timescales.

## INTRODUCTION

All cells face the fundamental challenge of providing a constant supply of energy, in the context of a rapidly shifting nutritional environment. The budding yeast *Saccharomyces cerevisiae* grows rapidly on glucose using “aerobic fermentation”, with ATP production almost exclusively from glycolysis (reviewed in ^1^). Preferential use of aerobic fermentation to quickly generate ATP occurs across diverse systems, including human skeletal muscles, activated T-cells and tumors, where it is termed the “Warburg effect” ^2,3^.

In response to glucose depletion, yeast and other organisms undergo “diauxic shift”. This entails a comprehensive transcriptional re-programming, in which hundreds of genes required for stress-responses and utilization of alternative carbon sources are upregulated ^4,5^. Simultaneously, expression of ribosome maturation and protein synthesis factors is suppressed. However, at nutrient shift the cellular mRNA pool is composed of pre-existing transcripts produced under glucose-replete conditions. Reprogramming is therefore complemented by more rapid post-transcriptional changes to acutely modify the functional proteome ^6–8^. Notably, global mRNA translation is dramatically reduced within minutes ^9,10^This is selectively bypassed for specific stress-induced transcripts ^11,12^. Together, this allows yeast to mount protective responses and dynamically alter metabolism immediately following glucose withdrawal.

Translation repression is a common response to numerous stresses, including temperature and osmotic shock, generally targeting initiation ^13,14^. Notably, translational shut-down following glucose withdrawal is distinct in both speed and scale ^8,15^. We previously reported that this results from a specific mechanism, in which key initiation factors are displaced from the 5’ regions of mRNA ^16^. Recruitment of the 40S and 60S ribosomal subunits at the start codon of a transcript is mediated by ‘scanning’ initiation factors, which unwind secondary structures in the mRNA and promote 43S-complex loading. These are eIF4B and two DEAD-box RNA helicases, eIF4A and Ded1^17–19^. Following glucose withdrawal, loss of binding by these factors blocks formation of 80S ribosomes. Translation activity ceases as elongating ribosomes reach the termination codons (see Fig1A).

Remarkably, glucose-dependent initiation factor release occurred within 30 sec, the earliest time point that could be tested ^16^. These proteins are extremely abundant; eIF4A is comparable in copies to the ribosome (∼150,000 per cell), while eIF4B and Ded1 also exceed 25,000 copies ^20^. The rapid regulation of such abundant targets presents a challenge: information on glucose depletion must be sensed, transmitted and amplified to alter protein binding within seconds. While rapid regulation of RNA-protein interactions mediates post-transcriptional responses to many cellular stresses, this is an unprecedented timescale of response ^21^. Here, we set out to investigate the mechanisms allowing yeast cells to achieve seconds-level remodeling of RNA metabolism following glucose withdrawal.

Although implicated in the response of yeast cells to multiple stresses, depleting intracellular pH did not recapitulate the effects of glucose depletion ^16^. Considering other metabolite regulators, we noted that the DEAD-box helicase initiation factors are ATP-dependent RNA binding proteins^22,23^. This requirement for ATP means their binding activity is subject to kinetic effects, alongside control by protein-protein interactions ^24^. Hypothesizing glucose-dependent changes, we discovered that rapid remodeling of NTP levels drives translation shutdown. This is a distinct mechanism from the major well-characterized stress-signaling pathways, and can be triggered by energy depletion without carbon source shift. The resulting translation shutdown is global, prior to preferential restoration on newly synthesized mRNAs. We conclude a biphasic adaptive response to energy stress: Metabolic remodeling, acting in seconds, mediates immediate protective changes in RNA metabolism, with subsequent transcriptional regulation by kinase/phosphatase cascades acting over minutes.

## RESULTS

### Rapid NTP depletion and initiation factor displacement

The translational response to glucose withdrawal is remarkably rapid, with global initiation shutdown achieved on a timescale of seconds. This appeared to be excessively fast for regulation by conventional stress-responsive signaling cascades. Contrastingly, levels of nucleotide triphosphates (NTPs) have been reported to change rapidly following loss of glucose in *S. cerevisiae* ^25,26^. The DEAD-box helicases are predicted to require ATP to support translation initiation, suggesting the possibility that an abrupt drop in ATP levels and/or ATP:ADP ratios might be responsible for their displacement from mRNA 5’ UTRs. In contrast, the major translation elongation factors utilize GTP. We hypothesized that the distinction in NTP utilization could underpin a mechanism for translation regulation acting on timescales shorter than most previous analyses.

We established rapid protocols to characterize changes in metabolites and RNA metabolism on a timescale of seconds. Nutrient shift and harvesting of exponential yeast cultures followed by metabolic quenching was achieved within 10 sec total time, or UV crosslinking within 30 sec (Fig. S1A). NTP and NDP levels were quantified using targeted HPLC separation coupled with UV detection ^27,28^. Quantified NTP recoveries were used to estimate intracellular nucleotide concentrations (see Methods). Rapid time courses over the first minute following glucose withdrawal were performed in biological triplicate and compared to mock shifts (Fig. S1B). The results were highly reproducible.

We initially applied this approach to a switch from medium containing 2% glucose (Glu), to medium lacking glucose but containing 2% glycerol plus 2% ethanol (Glyc/EthOH) to mimic diauxic shift. The measured intracellular ATP level was close to 2mM during growth on glucose medium. Following transfer to Glyc/EthOH medium, ATP dropped ∼7-fold to below 0.3 mM by 30 sec, remaining at this level through the 60 sec timepoint (Figs.1B, S1B; Table S2). Contrastingly, the cellular ADP concentration in glucose medium was 0.18 mM, but rose to 0.45 mM by 30 sec after transfer to Glyc/EthOH. In consequence, ADP rapidly became dominant over ATP (Fig. 1C, S2D; Table S2). These changes make it likely that many ATP-utilizing proteins are subject to both limiting substrate availability and product inhibition by ADP, within seconds of glucose withdrawal.

**Figure 1.**
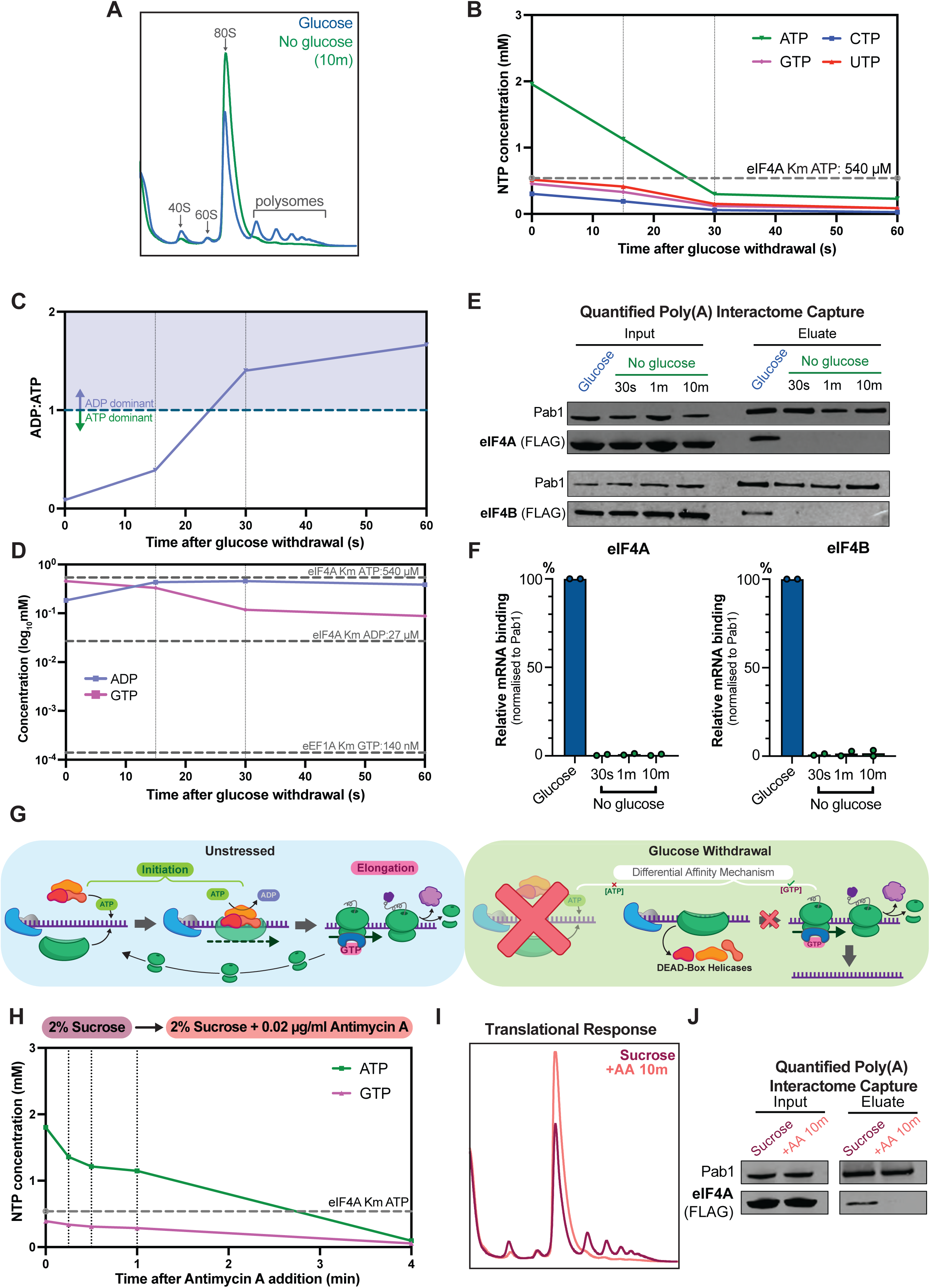
Rapid intracellular ATP depletion directly drives translation shutdown. A: Polysome gradient analyses assessing translational status on 2% glucose, and 10 min after transfer to 2% glycerol/ 2% ethanol (No glucose). Positions of small (40S) and large (60S) ribosomal subunits, monosomes (80S) and polysomes with multiple ribosomes are indicated. B: Estimated intracellular nucleoside triphosphate (NTP) pools following glucose withdrawal. (mean values; n=3). Km_[ATP]_ for eIF4A is indicated (dashed line). C: ADP:ATP ratio following glucose withdrawal (mean values; n=3). D: GTP and ADP concentrations (log10-adjusted means; n=3). Binding constants for eIF4A and eEF1A are indicated (dashed lines). E: Analysis of translation initiation factor recovery following *in vivo* UV crosslinking (254nm) and poly(A) selection. Cells were irradiated during growth on 2% glucose or 30 sec, 1 min and 10 min after transfer to 2% glycerol/ 2% ethanol. Total proteins in cell lysates (Input) and proteins recovered after mRNA binding to oligo[dT]_25_ beads (Eluate) were analyzed by western blots decorated with anti-FLAG for FLAG-eIF4A and FLAG-eIF4B. F: Quantification of E. Intensities were normalized to Pab1 as positive control for input and pull-down efficiency (n=2). Bar graphs represent the amount of eIF4A/4B bound to mRNA as a percentage of pre-shift level. G: Proposed “Differential Affinity” mechanism for rapid translational arrest. In the absence of glucose, flux through the glycolytic pathway is stopped immediately and NTP production falls, while usage continues, leading to rapid depletion. ATP levels fall below the binding constants for the low affinity DEAD-box helicases required for initiation, displacing these ATP-dependent RNA-binding protein from transcripts. Comparatively, elongation factors are high affinity and retain GTP binding, allowing translation to continue from initiated ribosomes. In the absence of recycling, polysomes run off leaving free translation machinery and transcripts. H: Intracellular ATP and GTP levels (mean values; n=3) during growth in 2% sucrose, and following addition of mitochondrial inhibitor Antimycin A (AA) to 0.02µg/ml. Km_[ATP]_ for eIF4A is indicated (dashed line). I: Polysome gradient analyses for yeast grown in 2% sucrose and 10 min after addition of 0.02µg/ml AA. J: Western blot analysis of translation initiation factor mRNA-binding. *In vivo* UV crosslinking was performed during growth on 2% sucrose or 10 min after addition 0.02µg/ml AA, followed by poly(A) selection and analysis as in panel D.

Our original model for selective loss of translation initiation postulated that elongation may be favored by delayed depletion of GTP relative to ATP. However, GTP, CTP and UTP were each depleted on a comparable timescale to ATP (Fig. 1B). In particular, the starting GTP concentration of 0.46 mM fell to 89 µM by 30 sec of transfer to Glyc/EthOH (Fig. S1B; Table S2). We therefore considered whether there might be differences in the NTP binding constants for the translation initiation and elongation factors. Based on published data, eIF4A and Ded1 both show low affinity for ATP, with binding constants of 540µM and 300µM respectively ^29,30^. These values are consistent with other DEAD-box ATPases reported from yeast ^31,32^. In consequence, ATP depletion following glucose withdrawal causes intracellular concentration to fall well below the Km within 30 sec. Due to DEAD-box ATPases binding ATP and RNA cooperatively, this is expected to result in release from mRNAs. Furthermore, eIF4A displays affinity for ADP an order of magnitude greater than for ATP (Fig. 1D). ADP binding promotes an ‘open’ state with low affinity for RNA ^33,34^. The rapid dominance of ADP following loss of glucose will therefore block further mRNA binding.

In contrast, reported binding affinities for GTPase elongation factors are very much lower: for example 140 nM for eEF1A (Cavallius and Merrick, 1998). The measured GTP concentrations remain an order of magnitude above this value after transfer to Glyc/EthOH, so their continued function is expected after glucose withdrawal (Fig. 1D).

To confirm the timescale of initiation factor displacement from mRNA transcripts, we developed a poly(A) interactome capture approach. RNAs were UV crosslinked to bound proteins in cells grown in glucose or at timepoints immediately following transfer to Glc/EthOH medium. Poly(A)+ RNAs were purified from cell lysates using oligo[dT]_25_ beads. Binding of FLAG-tagged initiation factors was quantified by western blotting and Licor imaging, using Pab1 as positive control for input and pull-down efficiency (Figs. 1E, 1F) This revealed that eIF4A is lost from mRNA transcripts, reaching undetectable levels within 30 sec, consistent with NTP remodeling (Fig 1E). eIF4B is the binding partner of eIF4A, and showed equivalently reduced mRNA interactions over the same timescale. Recovery of eIF4B was much lower in non-crosslinked controls, and eliminated by competition with artificial poly(A) (Fig S1C), confirming binding specificity. We conclude that the entire 43S loading complex is displaced from total mRNA within 30 sec.

From these results, we propose that a ‘Differential Affinity’ mechanism drives translational arrest following glucose withdrawal. ATP depletion to below the binding constants for the DEAD-box helicase initiation factors (∼0.35-0.54mM) leads to reduced RNA affinity and directly blocks the translation initiation process. In contrast, GTP levels remain well above the very low (∼140nM) binding constants for elongation factors, supporting continued function. Together, this results in orderly translation shutdown by preventing 80S ribosome assembly while allowing polysome run-off (Fig 1G).

### ATP depletion induces translational arrest independent of carbon source shift

To investigate whether changes in ATP concentrations are sufficient to arrest initiation, we characterized the translational response to intracellular ATP depletion independent of carbon source shift. For this we utilized the mitochondrial inhibitor Antimycin A (AA), which blocks ATP generation through respiration. Although the BY4741 yeast strain growing in 2% sucrose shows highly similar glycolytic growth to on glucose (see below), there is an increased metabolic contribution from respiration when utilizing the disaccharide. Under these conditions, AA strongly reduced intracellular NTP pools without an external change in carbon source availability (Fig. 1H).

NTP depletion occurred on a minute time scale after AA addition, allowing comparison of the timescales of metabolic remodeling and translation arrest. Although NTP depletion after AA addition was slower than following glucose withdrawal, by 4 min intracellular ATP levels had fallen to 0.1 mM (Fig. 1I, S1D; Table S2). This is well below the binding constant of eIF4A. Polysome gradient analyses, performed after 10 min to allow time for ribosome runoff, demonstrate the loss of translation (Fig. 1H). Poly(A) interactome capture at the same time point confirmed the reduction in eIF4A binding to mRNA (Fig. 1J). Polysome profiling on aliquots of the same culture are consistent with reduced eIF4A binding to mRNAs driving polysome collapse (Fig. S1E).

These results support ATP-dependent binding as a direct regulator of translation initiation factor function.

### NTP remodeling is actively controlled

The glycolytic enzyme hexokinase 2 (Hxk2) has been implicated in regulating transcriptional and post-transcriptional responses to glucose availability ^10,36,37^. Consistent with previous data, a *hxk2*Δ strain showed resistance to rapid translation arrest following glucose depletion (Fig. 2A). In wild-type cells polysome collapse occurs within 10 min (Fig. 1A), whereas these were maintained at 10 min and 25 min in *hxk2*Δ strains, indicating ongoing translation initiation. We considered whether this might reflect reduced metabolic flux through the glycolytic pathway favoring respiratory metabolism ^10,36^. However, *hxk2*Δ strains showed a wildtype growth rate on glucose and (Fig. S2A) and were almost insensitive to AA, indicating little dependence on mitochondrial respiration (Fig. S2A). Yeast expresses two other hexose kinases (Hxk1 and Glk1), which are sufficient to maintain glycolytic flux.

**Figure 2.**
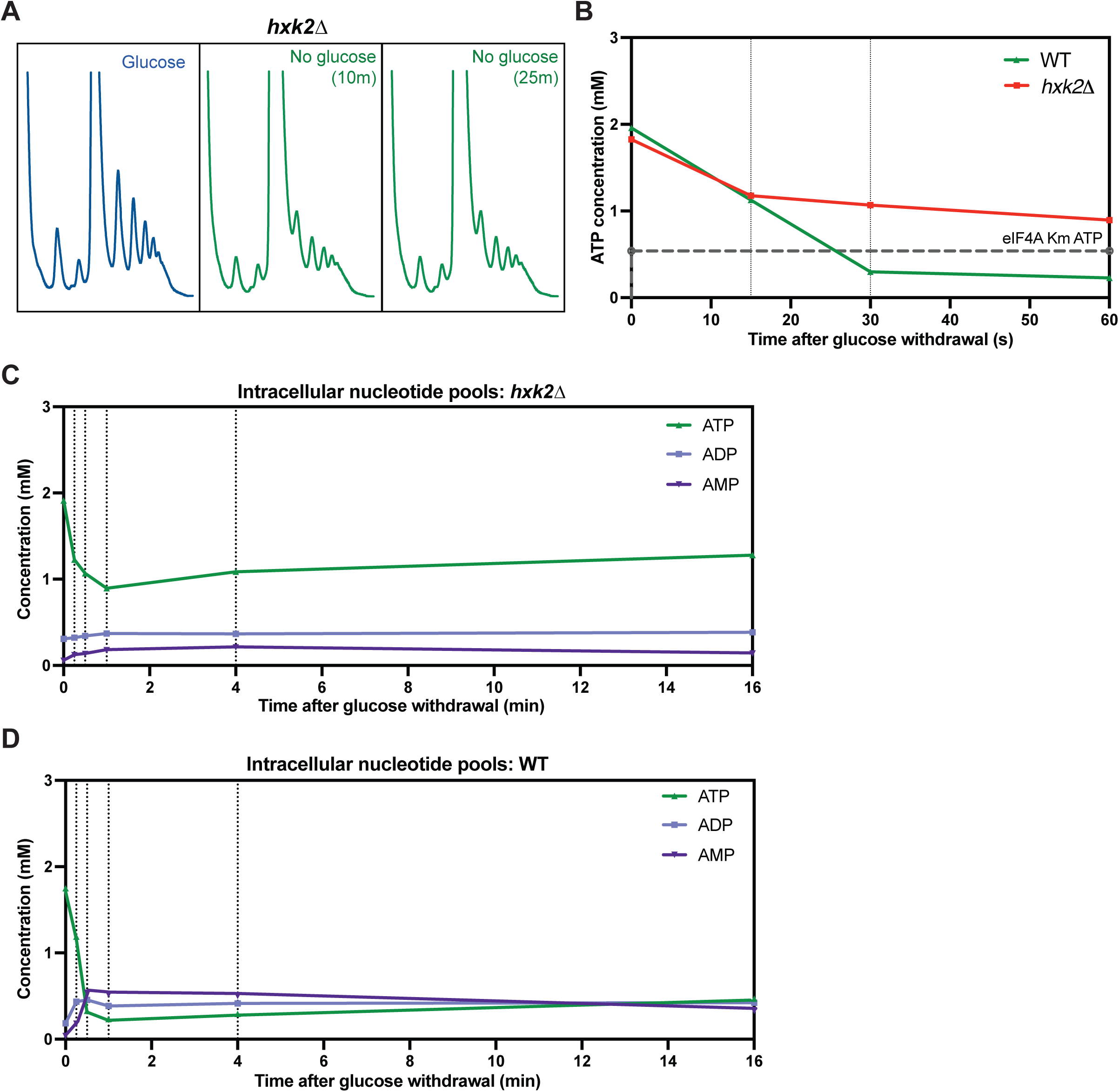
*hxk2*Δ mutants reveal active control of NTP remodeling. A: Polysome gradient analyses in *hxk2*Δ strains assessing translational status on 2% glucose, and 10 min or 25 min after transfer to 2% glycerol/ 2% ethanol (No glucose). B: Intracellular ATP concentrations (mean values; n=3) in wildtype and *hxk2*Δ strains following glucose withdrawal. C: Levels of adenosine nucleotides in *hxk2*Δ strain (mean values; n=6, except n=3 for 15 sec). D: Levels of adenosine nucleotides in wildtype (mean values; n=5, except n=2 for 15 sec).

Analyses of NTP levels indicate that continued translation in *hxk2*Δ mutants results instead from an altered metabolic response to glucose withdrawal (Figs. 2B, 2C, S2B-2D). The intracellular NTP concentrations decline initially (15 sec) comparably to wild-type cells (Fig. 2B). However, ATP levels then plateau at around 1mM, and this is maintained for at least 16 min. Concentrations of ADP and AMP also remain relatively stable in the *hxk2*Δ strain, as do the pools of CTP, UTP and GTP (Figs. 2C, S2B-2C). Since ATP levels remains well above the binding constants for the DEAD-box initiation factors, these results are consistent with ongoing translation initiation.

Notably, following initial depletion, the reduced ATP concentration also remains stable in wild-type strains over 16 min post-stress, as do ADP and AMP pools (Figs. 2D, S2D). Together, these results indicate that the depletion of NTPs, and dramatic alteration of adenylate nucleotide ratios, following glucose depletion is not simply an inevitable consequence of energy deprivation. Rather, a regulated response controls remodeling to achieve a new “set-point” for NTP levels.

### Rapidly growing yeast are primed for protective post-transcriptional responses

In previous analyses, the timescale and effects of glucose depletion were distinct from withdrawal form alternative sugars ^10^. These less readily metabolized sugars are expected to be predominately metabolized via mitochondrial respiration. We postulated that NTP remodeling on a second timescale occurs only in cells ‘primed’ by fast glycolytic growth to initiate a protective adaptation of metabolism under energy stress.

We used Antimycin A (AA) to assess relative dependence on oxidative phosphorylation. Yeast growing on raffinose displayed a lag period of 14 h, in response to very low doses of AA and failed to saturate (Fig 3A). Comparatively, yeast grown on sucrose demonstrated low AA sensitivity, only mildly greater than on glucose (Figs. 3B, S4A). Thus, sucrose and raffinose promote primarily glycolytic or slower respiratory growth, respectively. The doubling time on sucrose medium (∼97 min) is similar to glucose (∼98 min) but substantially faster than on raffinose (148 min) (Fig. S3A). Corresponding to these differences in metabolic state, we observe that transfer from raffinose medium did not cause precipitous NTP depletion (Figs. 3C, S3B), whereas shift from sucrose led to rapid loss of ATP equivalent to glucose withdrawal (Figs. 3D, S3C). Consistent with this, following raffinose withdrawal eIF4A was retained on mRNA over minute time courses (Fig. 3E). Polysomes were retained 10 min post-withdrawal, supporting ongoing translation initiation and protein synthesis. Conversely sucrose withdrawal caused loss of eIF4A binding to mRNA in <1 min and polysome collapse (Fig. 3F).

**Figure 3.**
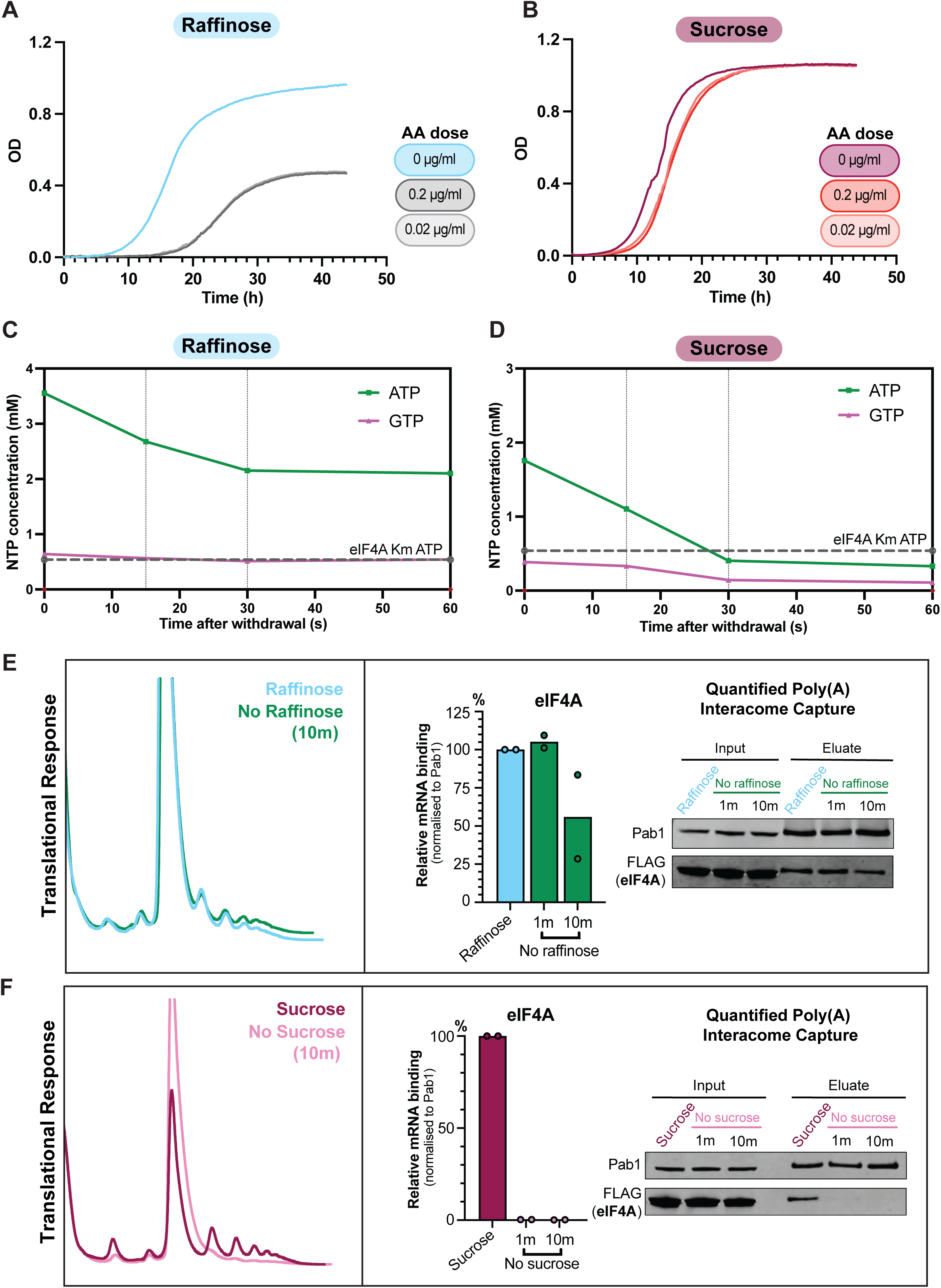
Metabolic changes are dependent on the prior carbon source. A: Growth curves (OD_600nm_) with raffinose as carbon source, in the presence of the mitochondrial inhibitor Antimycin A (AA) at 0.2µg/ml, 0.02µg/ml (n=2). B: As A, for yeast grown on sucrose. C: Intracellular NTP levels (mean values; n=3) following shift from 2% raffinose to 2% glycerol/ 2% ethanol. Km_[ATP]_ for eIF4A is indicated (dashed line). D: As C, for carbon source shift from 2% sucrose to 2% glycerol/ 2% ethanol. E: Translational response to raffinose withdrawal. mRNA association of eIF4A was assessed using poly(A)-interactome capture as described for Figures 1D and 1E. Bar graphs show eIF4A binding relative to the pre-shift level (n=2). Polysome profiles 10 min after raffinose withdrawal. F: As in E, for sucrose withdrawal.

We conclude the regulated NTP remodeling response to carbon source withdrawal is not glucose-specific, but a feature of fast-growing yeast. The metabolic response driving immediate, post-transcriptional changes appear specific for yeast growing rapidly by glycolysis.

### Energy depletion drives transcriptional reprogramming

Following glucose withdrawal, selective production of adaptive proteins depends on coordinated translational and transcriptional responses. To assess whether rapid NTP remodeling might also contribute to transcriptional adaptation, the response to sucrose withdrawal was compared to well-characterized changes following glucose derepression. Sucrose depletion and RNA sequencing were performed as described for glucose withdrawal ^38^. Replicates showed excellent reproducibility (Fig. S3D), permitting direct comparison of the transcriptome during depletion of each carbon source (Fig S3E, Tables S3-4).

Yeast utilizing glucose and sucrose possessed highly similar transcriptomes, consistent with largely comparable metabolism and growth (Figs 4A, S4A-B). Differences in expression were predominantly moderate, and for lesser expressed transcripts. Global gene expression patterns altered dramatically following depletion of either carbon source with striking similarity; 90% of transcripts differentially expressed following glucose depletion showed a comparable response to loss of sucrose (Figs 4B-D). Gene ontology (GO) over-representation analysis of the most changed genes following either withdrawal showed enrichment for near-identical biological processes, suggesting comparable remodeling of metabolism and protein production (Fig. S4C).

**Figure 4.**
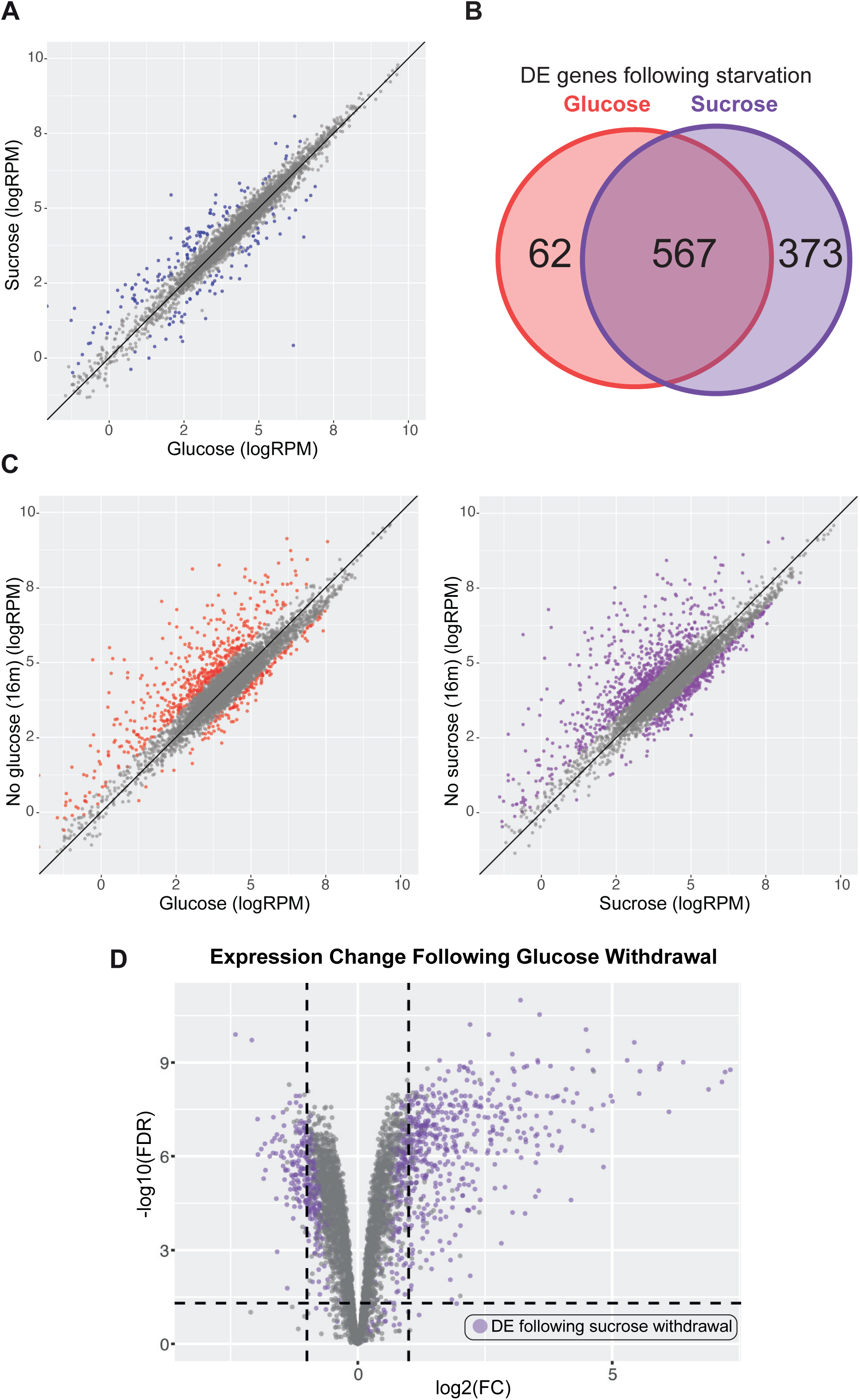
Transcriptional reprogramming in response to energy depletion. A: Transcript abundance (counts per million, RPM) in glucose versus sucrose media (n=3). Significantly altered mRNAs (fold-change >2, false-discovery rate <0.05) are colored blue. B: Venn diagram showing numbers of transcripts significantly altered following each carbon source withdrawal. C: Scatter plots comparing transcript abundance (RPM) following withdrawal of glucose (left panel) or sucrose (right panel) (n= 3 in each case). Significantly altered mRNAs (FC >2, FDR <0.05) are colored. D: Volcano plot showing the transcript level fold change distribution after glucose withdrawal for 16 min relative to control cells. For response comparison, transcripts differentially expressed (FC >2, FDR <0.05) following withdrawal of sucrose as carbon source are colored purple.

Yeast, and many other organisms, have glucose-specific signaling pathways (reviewed in ^39^). However, the similarities in transcriptional derepression between carbon sources indicate a more general response to energy deficiency. Direct effects of ATP depletion and/or ADP accumulation on regulators may be integrated with canonical signaling pathways.

### Immediate loss of translation initiation factors occurs globally

Diauxic shift requires widespread reprogramming of metabolism, and protein production stops while mRNA populations are modified. NTP depletion-driven loss of initiation factor binding is consistent with the inhibition of bulk translation following glucose withdrawal. However, translation is reported to continue or increase for important proteins that mediate stress responses and metabolic adaptation ^11,12^. We therefore considered whether these mRNAs specifically escape initiation factor eviction following glucose withdrawal, favoring their production under stress conditions. To assess this, we compared initiation factor binding to transcript abundance before and during glucose depletion (outlined in Fig. 5A).

**Figure 5.**
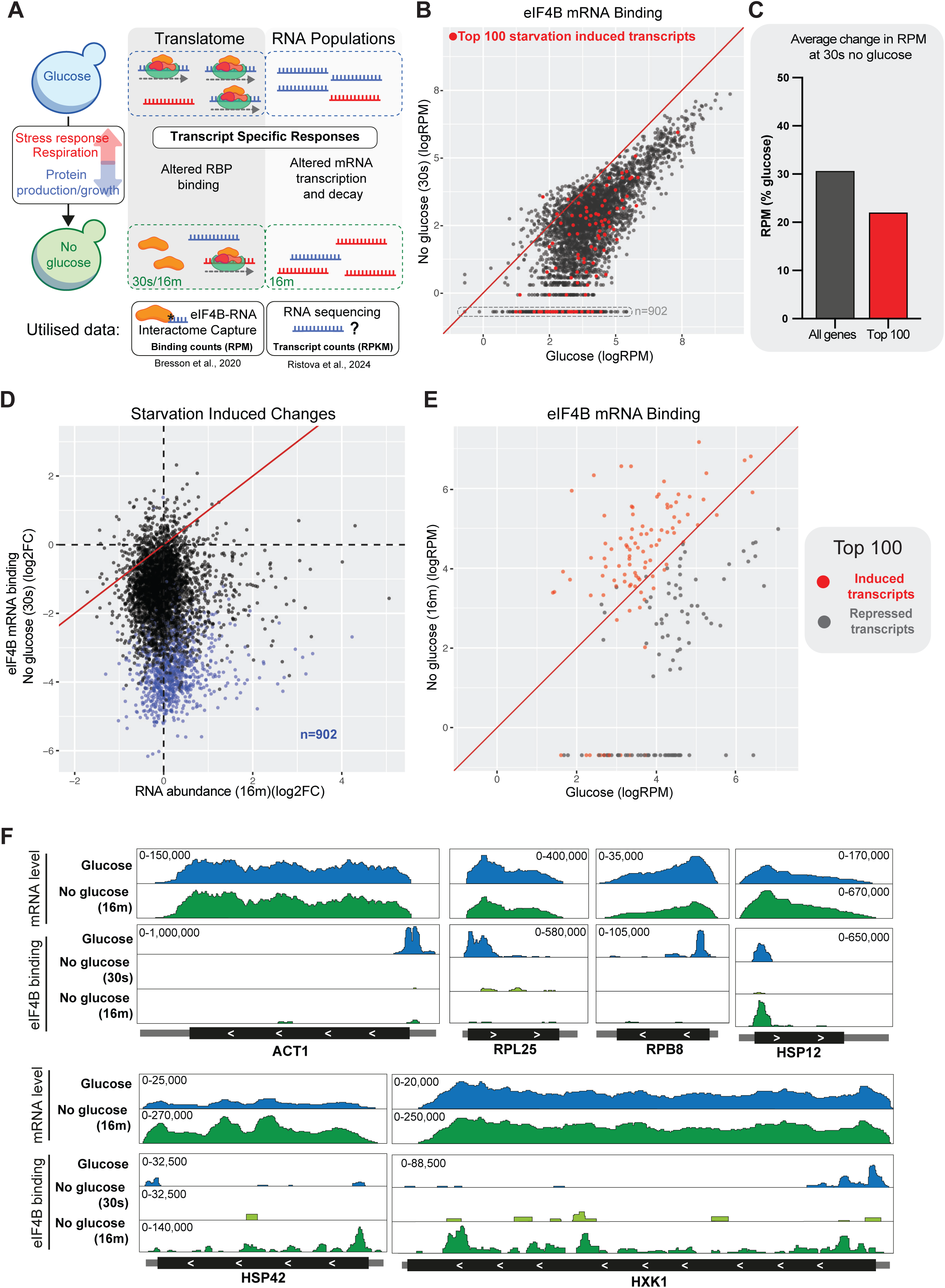
Immediate translation arrest occurs globally. A: Overview of bioinformatic approach to investigate transcript specific translational responses to glucose withdrawal. In response to glucose withdrawal, select transcripts are transcriptionally induced and translated against a background of global shut-down. To relate translation initiation factor displacement to such differential responses, transcript level data quantifying eIF4B binding prior-to or following glucose withdrawal (30 sec and16 min) was combined with time-matched RNA sequencing data. B: Scatter plot comparing changes in mRNA binding by eIF4B prior-to or following 30 sec glucose withdrawal. Transcript counts are normalized to library size, in Reads per Million (RPM). The 100 transcripts showing the greatest fold-change in abundance following 16 min glucose withdrawal (FDR <0.05) are colored red. For transcripts not detectably bound after glucose withdrawal, pseudocounts matching the minimum threshold (RPM=0.5) were added (lower, boxed line; n=902). C: Quantification of changes in B, (as % of RPM in glucose) for all detected (n=4,377) or most induced (n=100) mRNAs. D: Scatter plot comparing changes in eIF4B binding with subsequent changes in mRNA levels following energy stress. The fold-change (FC) in eIF4B association after 30 sec glucose withdrawal is compared to transcript abundance after 16 min. Undetected transcripts retained by pseudocount addition are indicated in blue; n=902. The x=y line is shown in red. E: Scatter plot comparing changes in eIF4B binding to mRNAs of the most differentially expressed genes following 16 min glucose withdrawal. The 100 transcripts with the greatest increase (red) or decrease (black) in abundance are shown (FDR < 0.05). mRNA binding by eIF4B is presented normalized to library size, in RPM. The lower line represents mRNAs maintained by pseudocount addition. F: Examples of eIF4B binding to “housekeeping” mRNAs, ACT1, RPL25, RPB8, or stress induced mRNAs, HSP12, HSP42, HXK1. Each set of tracks is normalized to library size (RPM), and relative scale is given in the top corner. Total mRNA abundance in RNA-seq is shown for comparison. For induced transcripts, scales differ between glucose and no glucose tracts. Open reading frames are indicated as black boxes (directionality indicated by white arrows), with UTRs as flanking gray boxes. Further examples are given in Fig. S5.

Poly(A)+ RNA sequencing data were obtained during growth on Glu medium or following transfer to Glyc/EthOH for 16 min ^38^. Abundance was quantified in sequence reads per kilobase of RNA per million mapped reads (RPKM), correcting counts for gene length and sequencing depth. Most mRNA abundances were little altered following glucose depletion, but stress-response transcripts and sugar transporters are already strongly upregulated by 16 min. Differential expression (DE) analysis identified the 100 most increased and decreased transcripts (FDR < 0.05), 16 min after glucose withdrawal (Tables. S4, S6).

As a proxy for initiation, we utilized data for RNA interactomes of eIF4A and eIF4B generated using UV-crosslinking (CRAC), which identifies bound RNAs and the location of the interaction site ^40^. Relative recoveries for each mRNA are quantified in sequence reads per million mapped reads (RPM). Crosslinking was performed during growth on Glu, and following transfer to Glyc/EthOH for 30 sec or 16 min (Fig 4A) ^16^.

Both eIF4A and eIF4B are lost from almost all transcripts following 30 sec in Glyc/EthOH medium, and this absence persists up to 16 min, consistent with poly(A) interactome capture (Figs. S5A, S5B). Note that binding to many mRNA species falls sufficiently to be undetectable by CRAC following glucose withdrawal (boxed points in Fig. 5B; blue points in Figs. 5D and S5A, S5B). To maintain these mRNAs in the data set, they were assigned “pseudo-counts” of 0.5 reads per million (RPM). For eIF4A, >50% of transcripts were not detected after stress. In consequence, we focused on eIF4B datasets for subsequent analyses.

We compared eIF4B binding to mRNAs on glucose and after transfer to Glyc/EthOH for 30 sec (no glucose) (Fig. 5B). Transcripts were filtered to produce a high confidence set (n =4,377), with reliable quantification (>0.5 RPM) in all replicates of at least one condition, removing mRNAs with very low expression (Table. S5). This analysis confirmed global displacement across almost all mRNA in seconds.

At 30 sec following glucose withdrawal, the 100 mRNAs that are most upregulated by 16 min showed no indication of preferential binding to eIF4B (Fig 5B, highlighted in red). Indeed, many lose binding sufficiently to fall below the detection threshold and appear in the imputed subset (lower horizontal line, n = 902), and eIF4B binding averaged for just this, subsequently induced, group is lower than for all mRNAs (Fig. 5C). Across the transcriptome, eIF4B binding immediately following glucose depletion (30 sec) shows no evident preference for mRNAs that are subsequently transcriptionally upregulated (Fig. 5D).

We also compared the location of eIF4B binding on individual mRNAs (Figs. 5F and S5C); including selected, abundant “housekeeping” mRNAs (shown for ACT1, RPL25 and RPB2), induced, stress-responsive mRNAs (HSP12, HSP42, HSP82, HSP104, HSP26) and upregulated glycolytic enzymes (HXK1, GSY1). On Glucose, most mRNAs show a clear peak of eIF4B binding around the AUG, as expected. At 30 sec after glucose depletion, eIF4B binding has been almost entirely lost on all mRNAs. At 16 min the housekeeping mRNAs remain very abundant, but show little restoration of eIF4B binding. In marked contrast, the induced genes show substantially increased eIF4B binding. However, alongside the 5’ peak there was increased binding throughout the body of the transcript. These strongly upregulated mRNA populations must largely consist of RNAs synthesized post-shift, whereas housekeeping transcripts will have been predominantly produced prior to this.

We postulated that the newly synthesized pool of each mRNA species has preferential access to the translation machinery. Supporting this, we compared the re-establishment of eIF4B binding on the sets of transcriptionally induced or repressed transcripts, defined by RNAseq (Fig 5E). 16 min after glucose depletion, strongly up-regulated transcripts nearly all showed increased eIF4B binding. In contrast, down-regulated transcripts predominately show reduced eIF4B binding; with 44% lost below detectable levels.

Preferential availability of newly transcribed mRNA species might reflect exclusion from condensates that sequester pre-existing mRNAs ^41^. As this new population is small, this potentially favors non-specific interactions with translation initiation factors, in addition to cognate 5’ binding.

### Newly synthesized mRNAs are preferentially translated

In principle, binding by translation initiation factors, or ribosomes, might not correlate with productive translation. We therefore assessed protein synthesis by metabolic labeling of nascent polypeptides using the “clickable” methionine analogue L-Azidohomoalanine (AHA). Labelling was performed for 16 min in glucose medium, or for 16 min immediately following transfer from glucose to glycerol/ethanol medium (Fig 6A). Total proteins were extracted, and newly synthesized proteins were purified by copper-catalyzed, covalent linkage to an alkyne-agarose resin. Following stringent washing, peptides were released by trypsin digestion and analyzed by mass spectrometry (MS). The total proteome was quantified in parallel and all replicates showed good correlation (Fig. S6). Unlabeled controls were included. Total proteomes (inputs) correlated well, but this correlation is lost following purification of AHA-labeled proteins supporting selective enrichment (Fig S7A).

**Figure 6.**
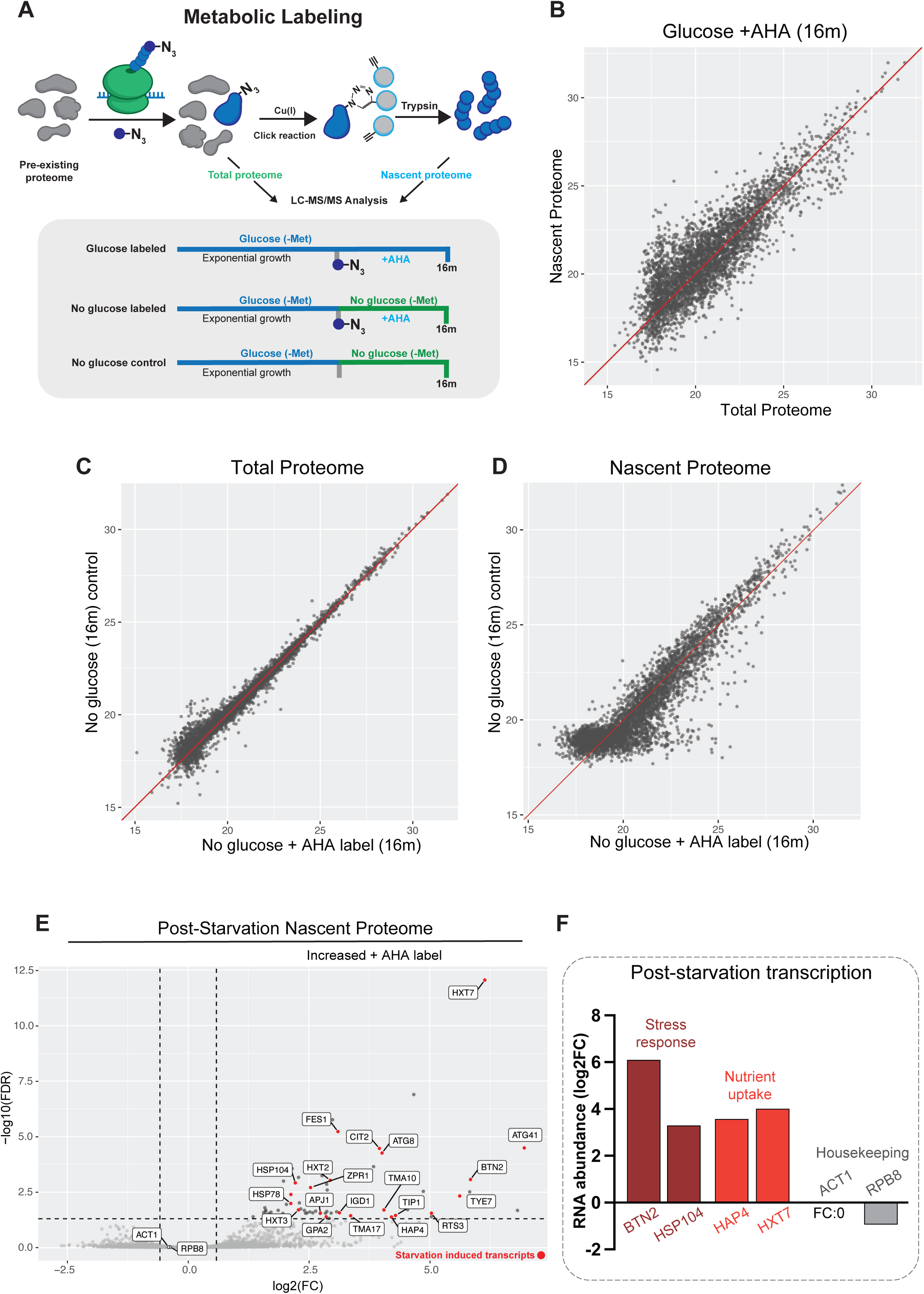
Metabolic labeling detects the newly synthetized translatome. A: Scheme of metabolic labelling experiment. L-Azidohomoalanine (AHA) is incorporated into nascent polypeptides. Cu(I)-catalyzed click chemistry covalently links the azide moiety in AHA to an alkyne-agarose resin, allowing stringent washing. Purified peptides are released by Trypsin digestion and analyzed by mass spectrometry (MS). Labeling was for 16 min during growth in 2% glucose, or commencing immediately after shift to 2% glycerol/ethanol (No glucose). Unlabeled controls were prepared in parallel. B: Scatter plot comparing protein intensity (log2) in the total and AHA-labeled proteomes during steady-state growth on glucose (n=3). C: Scatter plot comparing total proteomes (input) for AHA-labeled and unlabeled control samples following 16m glucose withdrawal. Median protein intensities (n=3) are plotted on log2 scale. D: Scatter plot comparing purified proteomes (nascent) from AHA-labeled and unlabeled control samples following 16 min glucose withdrawal. E: Volcano plot showing the fold change (FC) in purified proteomes (nascent) between AHA-labeled and control samples prepared 16 min following glucose withdrawal. Proteins with FC >1.5 and FDR < 0.05 were considered significantly differentially enriched. Enriched proteins whose mRNA levels also show significant increase (FDR <0.05) during glucose withdrawal are colored red and labeled. F: Changes in mRNA abundances for selected proteins involved in stress response and sugar uptake, compared to mRNAs for housekeeping proteins (highlighted in E).

During steady-state growth on glucose medium, the newly synthesized and total proteomes were in broad agreement, as expected (Fig. 6B). Following transfer to glyc/EthOH for 16 min, the control and labeled input samples were closely correlated (Fig. 6C), while there were clear differences following purification of labeled proteins (Fig. 6D). Differential proteomic expression analysis identified relatively enriched proteins following AHA-labeling, while no proteins were significantly enriched in the unlabeled control (Fig 6E). The most induced proteins are stress response factors and hexose transporters (labeled in Fig. 6E). Many enriched proteins are products of the 100 most induced mRNAs (highlighted in red in Fig. 6E). Selected, corresponding mRNAs are shown (Fig. 6F).

Globally, at 16 min most proteins are largely unchanged (Fig. S7B), but a set of mRNAs are induced for both transcription and nascent protein production (Fig. S7C). Differential expression analysis identified proteins with significantly different label incorporation prior to and following glucose withdrawal (n=524). Distinct groups of proteins showed higher label incorporation on glucose (cluster 2) or following glucose withdrawal (cluster 3), supporting preferential production under these conditions (Fig. 7A).

**Figure 7.**
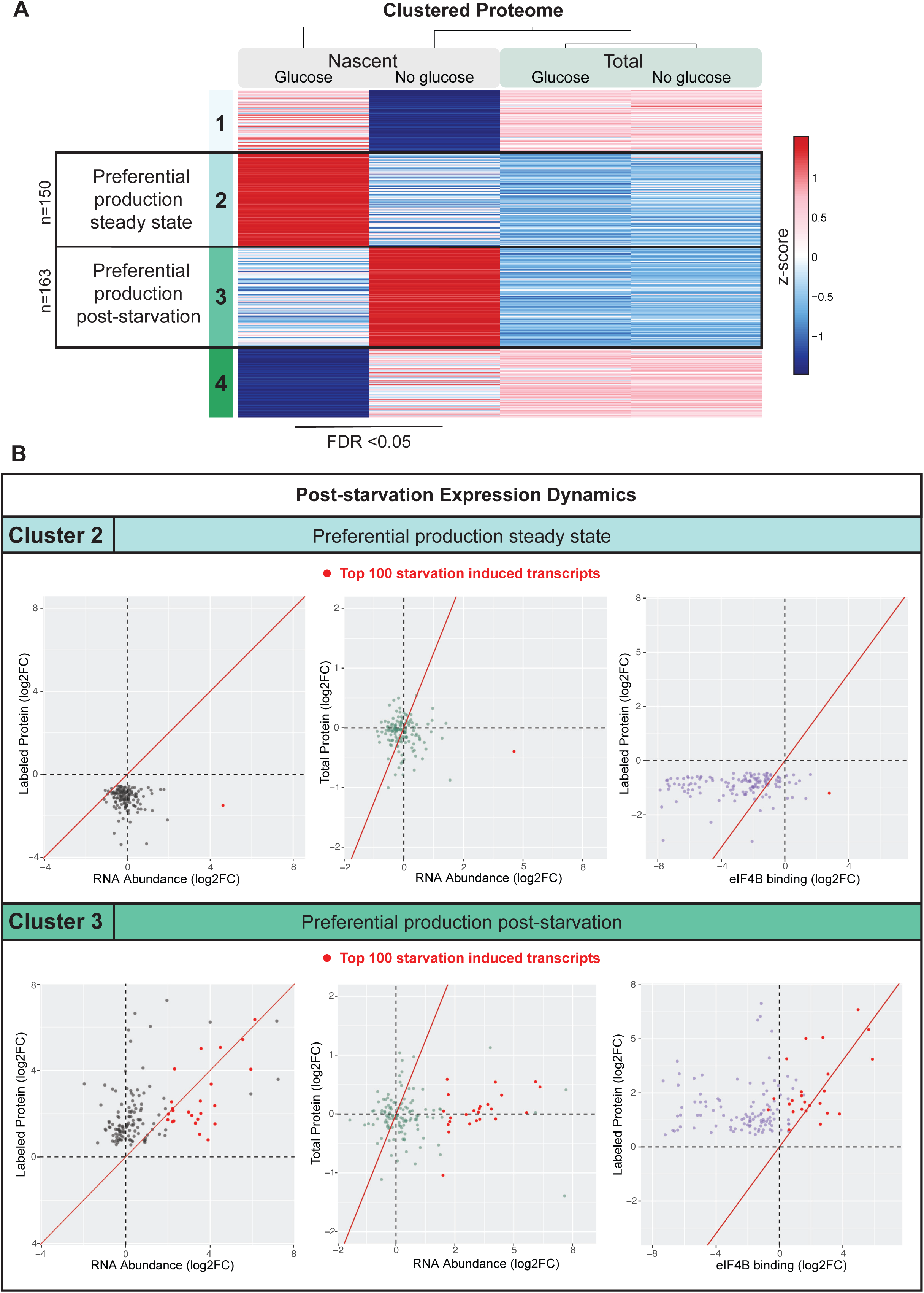
Newly synthesized mRNAs are preferentially translated. A: Cluster analysis of nascent protein production. 524 proteins exhibited significantly (FDR <0.05) different abundance in the proteome labelled over 16 minutes prior-to or immediately following glucose withdrawal. For the proteins, intensities in total and nascent proteomes were k-means clustered. B: Scatter plots for proteins in Cluster 2 (higher label incorporation in glucose) and Cluster 3 (higher label incorporation after glucose withdrawal). Graphs show fold change (FC) in mRNA levels (RPKM), eIF4B binding (RPM) and protein abundance (intensity) in glucose or following glucose withdrawal. The 100 proteins with most increased mRNA abundance following 16 min glucose withdrawal are colored red. The x=y line is shown in red.

Cluster 2, with higher translation on glucose was enriched for housekeeping functions. These show stable mRNA levels, low protein production and low association with eIF4B binding following glucose withdrawal (Fig. 7B). In contrast, cluster 3 includes 25% of the 100 mRNAs identified by DE as most upregulated following 16m glucose (Fig. 7C, highlighted in red in scatter plots), which correlates with increased protein production and elevated binding of eIF4B.

We conclude that eIF4B binding is initially heavily reduced for nearly all mRNA and contributes to attenuated translation. Subsequently, eIF4B preferentially binds newly produced mRNA. For a select, heavily induced population produced near entirely post-stress, this results in significantly increased protein translation. This sequence-independent mechanism promotes selective production of adaptive proteins following energy stress, dependent on temporal co-ordination of translational and transcriptional responses.

## DISCUSSION

Here, we report that that the immediate protective response to energy stress is driven by a specialized metabolic mechanism, distinct from kinase networks mediating conventional glucose repression.

Translational arrest following glucose withdrawal occurs through displacement of translation initiation factors, including DEAD-box helicases eIF4A and Ded1 ^16^. Strikingly, mRNA binding was abrogated within 30 sec of glucose depletion, a speed of response seemingly incompatible with conventional stress-responsive signaling cascades. DEAD-box helicases bind RNA and ATP cooperatively, whereas ADP binding promotes an open, non-RNA bound conformation ^33,34,42^. This suggested that altered ATP and/or ADP levels as glycolytic flux falls, might drive translation factor release.

Analyses of nucleotide availability revealed that intracellular ATP levels drop from ∼2 mM to below 0.3 mM within 30 sec following glucose depletion. This concentration is below reported ATP binding constants for DEAD-box helicases, including eIF4A (∼0.35-0.54 mM) ^29,30,34^. Over the same period, ADP rises from around 0.18 mM to 0.45 mM. This is substantially greater than the ATP concentration, or the reported eIF4A Km for ADP (0.027mM) ^33^, which favors an open structure with low RNA affinity. We propose that loss of ATP binding and increased ADP association directly drive release of eIF4A from all mRNA transcripts, blocking the initiation process. However, even after depletion, a minimal ATP-associated population will remain, potentially supporting the residual level of protein synthesis.

Over the initial period of glucose depletion, translation elongation is more resistant to inhibition than is initiation, resulting in bulk polysome run-off (Fig 1A) ^10,11^. Notably, translation elongation is more dependent on GTP than ATP. In our metabolite analyses GTP, was depleted with similar kinetics to ATP, presumably reflecting the requirement for ATP in GTP recycling. However, GTP levels remained substantially above the binding constants for elongation GTPases ^35^. Strikingly, these show GTP-binding affinities around 3,000-fold greater than the initiation factor ATPases. We therefore proposed a mechanism based on differential affinity. NTP depletion rapidly prevents 80S ribosome assembly but allows initiated ribosomes to continue and terminate, resulting in an orderly translation shutdown.

During termination, ribosome release is mediated by eRF1-eRF3, or the homologs Dom34-Hbs1, which are GTP-dependent (reviewed in ^43^). Peptide release by the ATPase Ril1 is independent of ATP hydrolysis, whereas 40S-60S splitting by Rli1 is ATP dependent ^44^. This may explain why ribosomes released from polysomes accumulate as 80S monosomes, rather than free subunits.

The metabolic response reflects energy depletion, rather than being glucose specific. Depletion of ATP by inhibition of respiration without carbon source shift, also resulted in the loss of initiation factor binding and ribosome run-off, supporting a direct ATP-driven mechanism. Similarly, withdrawal of sucrose caused rapid ATP-depletion and translation factor loss. Sucrose supports fast growth and is largely metabolized by glycolysis. In contrast, raffinose supports slow growth and is largely metabolized by respiration. Withdrawal of raffinose did not result in substantial ATP depletion or translation factor loss. We conclude that rapid NTP remodeling is ‘primed’ by fast glycolytic growth. Rapid ATP generation through glycolysis, with a minimal mitochondrial network, is highly efficient but results in vulnerability to energy stress ^45,46^. NTP remodeling may “de-risk” glycolytic growth by curtailing energy-consuming processes on very rapid timescales.

The differential affinity model requires that both enzymatic constants and NTP levels are “tuned” by evolutionary selection to enable distinct responses following glucose withdrawal. We noted that after rapid depletion during the first minute, NTP levels were approximately constant over the following 16 min. This suggested that the cells do not simply exhaust NTPs. Many cellular processes utilize NTPs, with a wide range of binding constants and turnover rates. We suggest that as the effective [NTP] drops, different NTP sinks will be progressively inhibited until residual consumption and production match, and a new equilibrium is established. In practice this occurs at ∼0.3 mM ATP after 30 sec. Residual ATP production may reflect a low level of respiration, or glycolysis from mobilization of stored glycogen.

In yeast lacking the glycolytic enzyme Hxk2, polysomes are not lost following glucose withdrawal ^10,36^. In *hxk2*Δ yeast, ATP levels initially declined but reached equilibrium faster and at a higher set point: 1.0 mM after 15 sec; above the binding constants of the initiation factors. We conclude that regulated metabolic remodeling controls NTP levels following glucose depletion. This higher set point is substantially above the binding constants of the initiation factors, suggesting that one or more major ATP sink is inactivated at a higher [ATP] in the mutant. A plausible candidate is the plasma membrane H^+^-ATPase (Pma1), which alone consumes around 30% of the ATP budget in WT cells and has been functionally linked to hexokinase in numerous studies ^47–50^.

Notably, subsequent changes in global gene expression patterns were strikingly similar following either glucose or sucrose withdrawal. 90% of transcripts responded comparably, reflecting remodeling of central carbon metabolism and repression of protein synthesis components ^4,5^. We postulate that a general “energy stress” program also initiates transcriptional reprogramming, independent of the canonical glucose repression network. We anticipate that subsequent metabolic remodeling allows respiratory metabolism, full restoration of ATPase function, and resumption of growth, after an extensive lag.

A range of environmental insults trigger translation repression. This inactivates pre-stress mRNAs while a selective transcriptional response is mounted ^51^. Importantly, while bulk synthesis is rapidly lost, stress-induced proteins must be selectively produced. Structural or sequence elements allowing specific evasion of translation repression have been suggested ^52,53^, but widespread common features have not been found. Rather, reporter systems indicate that timing is crucial; mRNAs transcribed after the initial stress escape repression ^41,54^. Supporting a sequence-independent mechanism for selective translation, our analyses found no evidence that withdrawal-induced mRNAs escape initiation factor eviction following glucose withdrawal. Instead, eIF4B was globally displaced on a second timescale, and then re-engaged over minutes preferentially on transcriptionally upregulated transcripts. Concordantly, initiation factor binding patterns shifted on these transcripts to reveal non-specific interactions alongside cognate 5’ binding. This indicates reduced specificity, possibly reflecting an excess of initiation factors relative to available mRNAs.

Immediately following glucose withdrawal, release of the protein-synthesis machinery leads to sequestration of free mRNAs into cytoplasmic condensates. However, newly transcribed mRNAs are excluded from condensates, by an unknown mechanism ^54^. Reduced global translation will generate free pools of translation initiation factors but these can access only the available, newly synthesized mRNA pool. A relative excess of initiation factors may favor translation even under conditions of ATP limitation.

Ded1 and other DEAD-box helicases participate in mRNA condensate formation under stress conditions. However, condensates form via multivalent interactions and the contribution of RNA binding in the helicase active site is unclear. Ded1 is lost from the AUG-proximal regions of all mRNAs immediately after glucose withdrawal, but remains associated with some coding sequences, potentially in condensates ^16^. This indicates that alternative modes of interaction exist and can be maintained after ATP depletion, perhaps facilitating roles in condensate formation and other regulatory processes. We propose that evolutionary tuning of enzymatic constants to control NTP remodeling allows specific rapid modulation of RNA metabolism following carbon-source shift.

Our findings may be relevant to other systems. ATP abundance in glucose was 1.7 x 10^-16^ mol per cell, corresponding to approximately 10^8^ ATP molecules per cell. After glucose withdrawal, the ATP level drops by ∼8x 10^7^ molecules in 30 sec, giving an ATP depletion rate of 3x 10^6^ molecules per second in yeast. During normal metabolism in human cells, the ATP pool is reported to be around 10^9^ molecules, and is entirely turned over in around 1 – 2 min ^55,56^. Therefore, a complete block in ATP production should result in a comparable ATP depletion rate to that seen in yeast. Moreover, 70-80% ATP depletion is reported for focal ischemia following arterial occlusion (see ^57^ and references therein), or with combined oxygen and glucose deprivation in neuronal cultures ^58,59^.

NTPases function in a vast range of essential and non-essential activities. Selection for differential affinity may provide a single mechanism to coordinate comprehensive post-transcriptional adaptation to energy stress. Similar enzymatic tuning might have modulated binding constants for many metabolites, such that alterations in metabolite abundance are buffered by hierarchical loss of selected pathways.

## LIMITATIONS OF THE STUDY

Loss of eIF4A and eIF4B is expected to block translation initiation, but other initiation factors may also contribute to inhibition. It remains unclear how the ATP concentration “set point” is established and maintained as a new equilibrium following glucose withdrawal. It is unclear what features distinguish newly synthesized mRNAs from the pre-existing pool. These could be RNA structure and/or modification, associated proteins or localization – and these possibilities are not mutually exclusive. Finally, we do not yet know whether a similar strategy is applied in other systems as an immediate response to energy depletion or other stresses.

## ACKNOWLEDGEMENTS

We thank Stefan Bresson for his support in relation to this work, including the gift of cell lines, consultation regarding analysis and critical reading of this manuscript. Aleksandra Helwark also generously provided reagents, advice and support throughout. We thank Richard Clark from Clinical Research Facility Edinburgh for sequencing services; Cristina Cardenal Peralta for assistance with mass spectrometry; Shaun Webb for bioinformatics assistance and Nick Gilbert for generously providing training and access to facilities. KB was supported by Wellcome Trust PhD Studentship [218470]. MR was supported by an EASTBIO PhD fellowship [543KOM/G40389]. SS and AC were supported by the Swedish Cancer Society [22 2377 Pj] and Swedish Research Council [2022–00675_VR]. DT was supported by Wellcome Principal Research Fellowships [109916, 222516]. This work was facilitated by funding for the Wellcome Discovery Research Platform for Hidden Cell Biology [226791] and we gratefully acknowledge support from the Bioinformatics core. Work in the Wellcome Centre for Cell Biology was supported by Centre Core Grants [092076 and 203149].

## STAR METHODS

### Key resources table

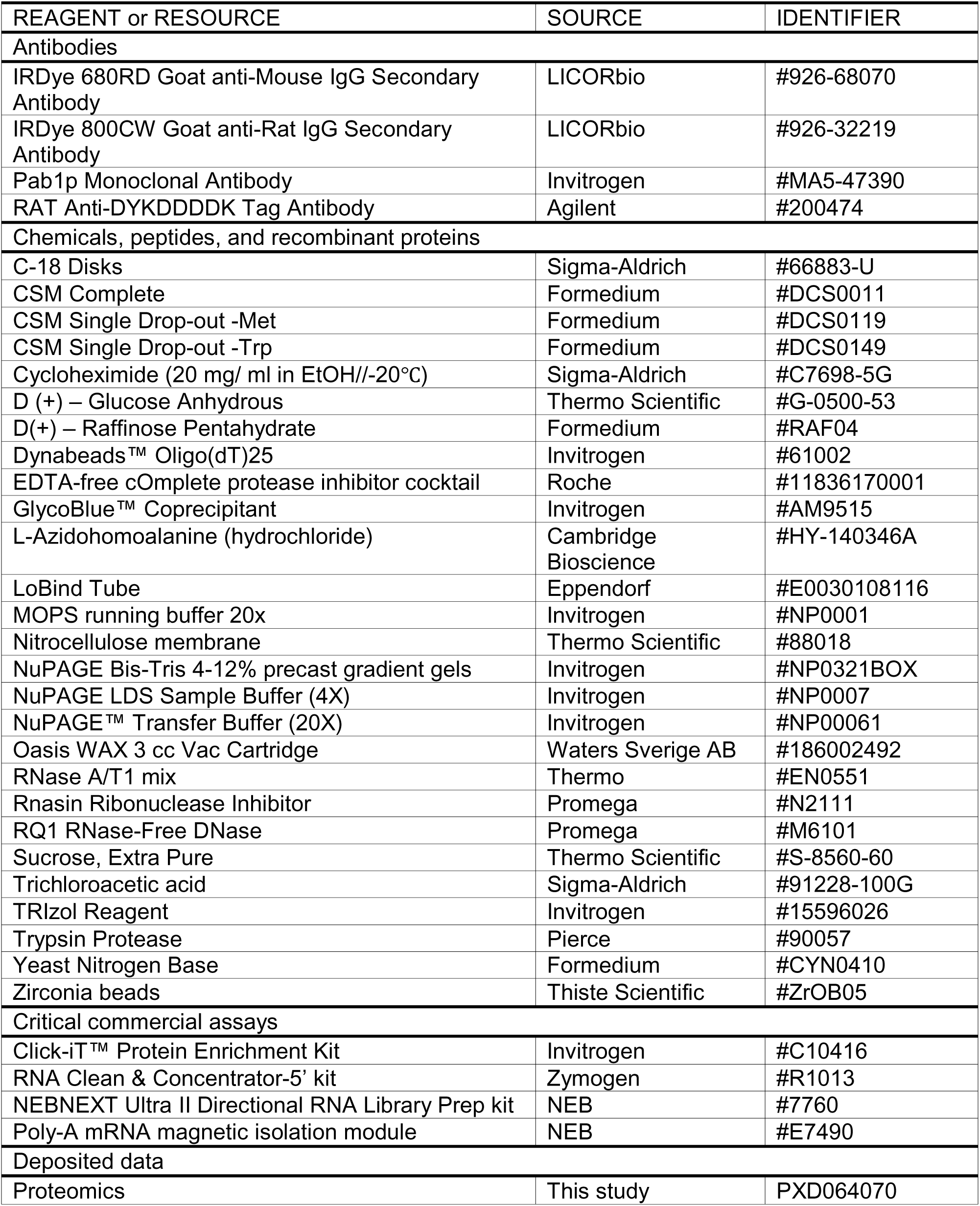
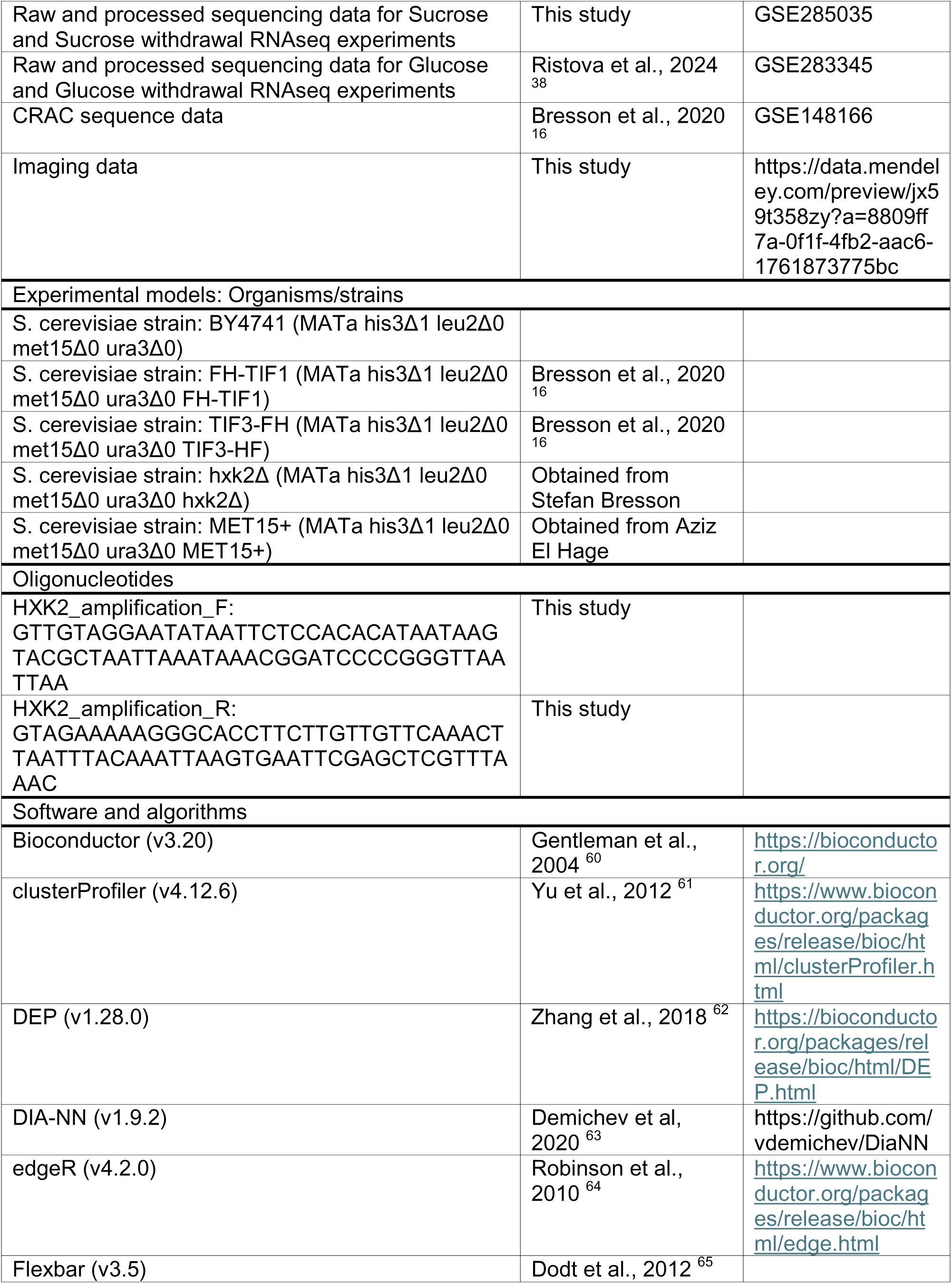
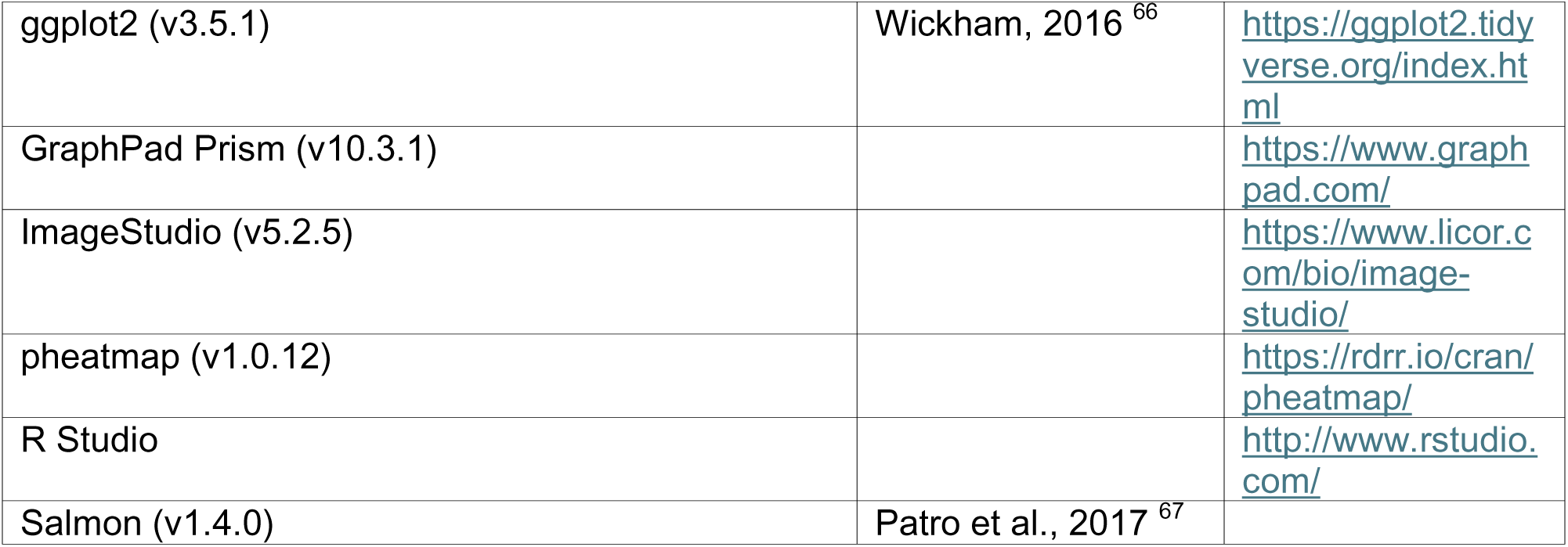

#### Reagents

All reagents used in this study are listed in the Key Resources Table.

#### Strains and Oligonucleotides

Strains generated or utilized in this study are listed in Key Resources Table and are available from the lead contact upon request.

#### Software and Algorithms

Bioinformatic tools utilized in these analyses are listed Key Resources Table.

#### Deposited data

The RNAseq raw data generated here, as well as utilized RNAseq and CRAC data sets reposited previously, are listed in Key Resources Table with relevant identifiers.

#### Experimental Model and Strain Construction

All S. cerevisiae strains used in this study were derived from the BY4741 background (*MATa his3*Δ*1 leu2*Δ*0 met15*Δ*0 ura3*Δ*0*). The methionine prototroph (BY4741 MET15+) carrying the WT S288C MET15+ gene, was utilized for metabolic labelling (generously gifted by Aziz El Hage). The *hxk2*Δ strain was a kind gift from Stefan Bresson. Conventional homologous recombination was used for complete deletion of the *HXK2* gene, utilizing the His3MX6 cassette from the pFa6a plasmid (Addgene #41596). Oligonucleotides were designed with 50bp homology to the *HXK2* locus, detailed in Key Resources Table. All tagged strains were constructed and utilized previously in Bresson et al., 2020, as described. The chromosomal copy of TIF1 (one of two genes encoding eIF4A) was N-terminally tagged with Flag-His (FH), consisting of a single Flag motif, a four-alanine spacer, and eight consecutive histidine residues (DYKDDDDKAAAAHHHHHHHH). TIF3 (eIF4B) was C-terminally tagged with the same elements in reverse (HHHHHHHHAAAADYKDDDDK), the His-Flag (HF) tag. Both strains were generated using CRISPR-Cas9 ^68^.

#### Cell culture and medium

Yeast strains were cultured at 30°C with constant shaking at 200 rpm in 2% synthetic complete (SC) 2% yeast nitrogen base medium supplemented with 2% carbon source, as indicated. For all experiments, saturated overnight starter cultures were inoculated into fresh media at a starting OD_600_ of 0.05 and grown to exponential OD (0.3-0.5). Glucose, sucrose and raffinose grown cells were harvested directly from supplemented media. For treatments, cells were collected by filtration on a nitrocellulose membrane and transferred to fresh medium. This medium contained the same carbon source (mock shift), 2% glycerol and 2% ethanol (carbon source withdrawal), or the same carbon source and either 0.2 µg/ml or 0.002 µg/ml Antimycin A (+AA), as indicated.

For poly(A)-interactome capture experiments, yeast cells were cultured in 2% synthetic dropout (SD) -TRP, supplemented with 2% yeast nitrogen base and 2% indicated carbon source. This minimizes potential interference during UV-crosslinking.

L-Azidohomoalanine (AHA) labelling experiments were performed with SD-MET media supplemented with 2% yeast nitrogen base and 2% indicated carbon source. Labelling was performed by addition of 200 µM AHA for 16 minutes, either directly to exponential glucose grown culture, or alongside carbon source withdrawal by transferred to glycerol and ethanol medium supplemented with AHA.

#### Growth Curves

For Antimycin A sensitivity assays, cultures were saturated overnight in SC medium supplemented with indicated carbon source. Subsequently, these were diluted to OD_600_ 0.05 in 20ml fresh media and grown to an OD_600_ of 0.3. Yeast cells were then shifted to SC medium supplemented with indicated carbon source and either 0, 0.2 or 0.02 µg/ml Antimycin A. In each case, 3 ml culture was pelleted at 4,000 rpm for 3 min, washed once and then resuspended in 3 ml shift medium. These cultures were diluted to an OD_600_ of 0.025 in the same media, and 150 µl aliquots plated into a 96-well plate in technical triplicate. Optical density was monitored over 48 h using a Sunrise TECAN plate reader, measuring at 15 min intervals with continuous shaking at 30°C. All conditions were tested in at least biological duplicate.

#### Polysome profiling

Overnight cultures of yeast were diluted to OD_600_ 0.05 in 100 mL SC medium supplemented with indicated carbon source, and cultured with shaking at 30°C. At OD_600_ 0.45, cultures were either harvested directly or collected by filtration and transferred to indicated treatment medium for 10 min. For *hxk2Δ*, 25m glucose withdrawal was also performed. In all cases, cells were then pelleted by centrifugation at 4,600 rpm for 2 minutes at 4°C, washed once with 20 ml of ice-cold water, snap-frozen in a dry-ice and ethanol bath, and stored at -80°C. Biological duplicates were collected for each condition.

The technique for lysis and profiling was modified from ^69^. Briefly, cells were resuspended in 300 μl of buffer (20 mM HEPES-KOH, 7.4; 100 mM KOAc; 2 mM MgOAc; 0.1 mg/ml Cyclohexamide; EDTA-free protease inhibitors). The suspension was transferred to 2 mL screw-cap tubes containing 200 μL zirconia beads and lysed using a Fast Prep-24 machine (MP Biomedicals) with a pre-cooled CoolPrep™ adapter. Four cycles were performed at 6 m/s for 40 sec, with cooling on ice in between rounds. The lysate was diluted with an additional 200 μl lysis buffer, and cleared twice by centrifugation at max speed for 5 min at 4°C.

350 μg of RNA was loaded onto 10%-45% sucrose gradients in 1X gradient buffer (10LmM Tris-acetate, pH 7.4; 70LmM NH_4_OAc; 4 mM MgOAc) prepared using the Gradient Master (BioComp) and refrigerated prior to use. Subsequently, the gradients were centrifuged in an SW40-Ti rotor in an Optima XPN-100 Ultracentrifuge (Beckman Coulter) at 38,000 rpm for 2.5 h at 4°C. Absorbance profiles were visualized using the Piston Gradient Fractionator (BioComp) equipped with a TRIAX flow cell (BioComp) for UV profiling. For comparative analysis, starting absorbances were zeroed and the monosome peaks then aligned.

#### Metabolomics

For each metabolomics experiment, BY4741 were grown in 150 ml SC media supplemented with indicated carbon source. At OD_600_ 0.3, exponential cultures were harvested by filtration and either lysed immediately or subject to the indicated treatment (Fig S1A). For this, 10 ml shift media was applied directly to the filter and incubated for the time indicated. With 10 sec remaining, shift media was removed using the vacuum and the filter rapidly transferred for lysis.

Where timepoints exceeded 1min, the filter was transferred to 150ml stress media and cultured with shaking at 30°C and re-filtered prior to the time-point. In all cases, cells were lysed by placing the filter in a petri dish containing 4 ml 12% Trichloroacetic acid (w/v) in 15 mM MgCl_2_, kept on ice. The lysis solution and filter were then transferred to a 5 ml Eppendorf tube, vortexed vigorously for 5 min and snap frozen in liquid nitrogen. These were stored at −80°C. All conditions were tested in at least biological triplicate, ensuring ∼ 600 million yeast cells were collected in each experiment.

For NTP quantification, thawed samples were vortexed for 30 sec then centrifuged at 4,000 rpm for 5 min at 4°C. 1 ml of the supernatant was transferred to a 1.5 ml Eppendorf tube and neutralized twice in a 1:1 ratio with a dichloromethane (DCM)–trioctylamine (TOC) mixture (1 ml DCM and 0.28 ml TOC). Following centrifugation at 14,000 rpm for 2 min at 4°C, 900 µl of the supernatant was transferred to new Eppendorf tubes. A 100 µl aliquot of the extracted nucleotide solution was then mixed with 1.15 ml Milli-Q water, and the pH was adjusted to between 3 and 4 using 0.5 µl 6 M HCl. Subsequently, analysis was performed by HPLC UV as described in ^27^.

The remaining 800 µl of extracted solution was used to quantify NDPs and AMP as in ^28^. Briefly, samples were adjusted to pH 4.6 and cleaned by OASIS WAX-SPE. Following elution, samples were dried using a Speedvac and resuspended in 200µl water. For HPLC analysis, 20 µl was loaded and run on a 4.6 mm × 150 mm Sunshell C18-WP 2.6 μm column (ChromaNik Technologies Inc.).

This methodology produces UV chromatograms for experimental samples accompanied by NTP, NDP and AMP standards. By comparison of the peak heights or areas, the amount of NTPs, NDPs and AMPs in yeast cell samples is calculated. The result is expressed as pmol/10^8^ cells. From these, we estimate intracellular nucleotide concentrations as in ^70^. Briefly, the volume of the soluble fraction of a haploid yeast cell was estimated as 45 x10^-12^ g (∼45 µm^3^) by subtracting the reported dry weight from wet weight (15 x10^-12^ g and 60 x10^-12^ g, respectively). Assuming a conversion of 1g/L, a volume of 4.5 x10^-14^ L/cell was used to transform quantified NTP levels into concentrations in mM, as reported in Supplementary Table 5.

#### Poly(A)-Interactome Capture

The poly(A)-interactome capture protocol is based on the experimental procedures in ^71^ and ^72^, with significant modifications.

For each experiment, Flag-tagged strains were grown in 800 ml SC -TRP media supplemented with indicated carbon source. At OD_600_ 0.4, cultures were either utilized directly or subject to the treatment indicated using the rapid shift protocol outlined in Supplementary Figure 1A. Here, the exponential culture was collected by filtration, transferred to 800 ml of relevant shift media, resuspended and incubated shaking until 20 sec prior to the time-point. All cultures were then UV-irradiated at 254 nm for 20 sec using the Vari-X-Link crosslinker ^73,74^. Following crosslinking, cells were collected by filtration and resuspended in 50 ml ice-cold H_2_O. Cells were then centrifuged at 4,600xg for 2 min, and pellets immediately stored at −80°C. All conditions were tested in biological duplicate.

For lysis, cell pellets were resuspended in 800 µl lysis buffer (100 mM Tris-HCl, pH 7.5; 500 mM LiCl; 10 mM EDTA; 1% Triton X-100; 5 mM DTT; 100 U/ml RNasin; complete EDTA-free protease-inhibitor cocktail). Following addition of 1.5 ml Zirconia beads to the suspension, cells were broken mechanically using six one-minute pulses on a benchtop vortex, with cooling on ice in between. The cooled lysate was centrifuged at 4,600xg for 5 min at 4 °C, and the supernatant was then transferred to Eppendorf tubes and cleared by centrifugation at 20,000xg for 20 min at 4 °C. This extract was diluted to 15 mg/ml in lysis buffer. For competed controls, 500 µl aliquots of extract were supplemented with 20 µl 10 mg/ml poly(A).

To capture polyadenylated RNAs, for each extract, 200 µl (∼1mg) of oligo(dT)_25_ Dynabeads were first equilibrated by four washes in lysis buffer (as above, except 10 U/ml RNasin). These were then incubated with 500 µl (∼7.5mg) extract for 40 min at room temperature, while rotating. Subsequently, using a magnet to capture the beads, the supernatant was recovered into a fresh tube from the beads for repeat incubations (described below). The beads were washed once with 500 μl wash buffer A (10 mM Tris-HCl, pH 7.5; 600 mM LiCl; 1 mM EDTA; 0.1% Triton X-100; 10 U/ml RNasin) and twice with 500 μl wash buffer B (10 mM Tris-HCl, pH 7.5; 600 mM LiCl; 1 mM EDTA; 10 U ml−1 RNasin). To elute RNA-protein complexes by temperature, the beads were re-suspended in 60 µl elution buffer (10 mM Tris-HCl, pH 7.5) and incubated at 80 °C for 2 min. The eluate was then collected immediately.

To fully deplete samples of poly(A) RNAs, this procedure was repeated twice further. The beads were retained, and re-equilibrated with four washes in lysis buffer before the preserved supernatant was re-applied. The three sequential eluates were then combined and treated with 4 μl RNase A/T1 mix in 20 µl RNase buffer (10 mM Tris-HCl, pH 7.5; 300 mM NaCl; 5mM EDTA, pH 7.5) for 1 h at 37°C. Recovered proteins were precipitated overnight at −20°C, using 2µl glycoblue in 1 ml acetone. These were pelleted at 20,000xg for 20 min, dried briefly and resuspended in 18 μl 1X NuPAGE LDS sample buffer supplemented with 50 mM DTT.

#### Western Blot Analysis

Western blot analysis was used to compare amounts of Flag-tagged initiation factors in inputs and eluates of poly(A) interactome capture. Poly(A)-binding protein (Pab1) acted as a control for input and pull-down efficiency.

Input samples were prepared by mixing 1µl 15mg/ml cell lysate with 1X NuPAGE LDS sample buffer supplemented with 50 mM DTT, and preparation of eluates is outlined above. All samples were boiled for 5 min, and resolved on a 4%–12% Bis-tris NuPAGE gel by electrophoresis at 150 V in 1X NuPAGE MOPS buffer. Proteins were transferred to nitrocellulose membrane using the Biorad Mini Trans-Blot apparatus with NuPAGE transfer buffer for 1 h at 100V. Membranes were blocked with 5% milk in PBS for 30 min, and probed overnight at 4°C with the indicated antibodies diluted in PBS with 0.1% Tween-20 (PBS-T)). Following washing in PBS-T, membranes were incubated in the corresponding fluorescently labelled secondary antibody for 1 h at room temperature. Membranes were subsequently washed again, then visualized by chemiluminescent imaging using the LiCor Odyssey CLx imaging system (LICORbio). Quantitative analysis of captured images was performed with the Image Studio software (LICORbio).

The following antibodies were used: rat Anti-DYKDDDDK Tag Antibody (1:1000; Agilent, 200474), mouse Pab1p Monoclonal Antibody (1G1) (1:1000; Invitrogen, MA5-47390), IRDye 800CW Goat anti-Rat IgG Secondary Antibody (1:5000; LICORbio, 926-32219), IRDye 680RD Goat anti-Mouse IgG Secondary Antibody (1:5000; LICORbio, 926-68070).

#### L-Azidohomoalanine (AHA) labelling and Click Chemistry

To assess post-starvation protein synthesis, metabolic labelling was performed methionine analog AHA, modified with an azide moiety to allow chemoselective ligation with an alkyne group. This “click " reaction allows efficient covalent capture of labelled nascent proteins and identification by mass spectrometry.

BY4741 MET15+ were grown to OD_600_ 0.4 in 200 ml SC -TRP media supplemented with 2% glucose. Cultures were then labelling for 16 min with AHA at final concentration 200µM, diluted from a 200 mM L-Azidohomoalanine (hydrochloride) (Cambridge Bioscience) stock dissolved in equimolar NaOH to neutralize pH. For glucose samples, this was added directly to the media, whereas starved samples were prepared by rapidly shifting (Supplementary Figure 1a) the exponential culture to 200 mL AHA-containing SC -TRP media supplemented with 2% glycerol and 2% ethanol. Labelled cultures were incubated shaking until the timepoint, collected by filtration and resuspended in 50 mL ice-cold H_2_O. Cells were pelleted at 4600xg for 2 min, snap-frozen in a dry-ice and ethanol bath, and pellets immediately stored at −80°C. Unlabeled controls were prepared simultaneously, and all conditions were tested in biological triplicate.

Total protein extraction and enrichment of newly synthesized proteins was performed using the Click-iT™ Protein Enrichment Kit (Invitrogen), with minor adaptations to the protocol. For lysis, cell pellets were resuspended in 800 µl lysis buffer (8 M urea, 200 mM Tris pH 8, 4% CHAPS, 1 M NaCl, supplemented with 2X complete EDTA-free protease-inhibitor cocktail. Cells were broken mechanically using a benchtop vortex, by addition of 1.5ml Zirconia beads to the suspension and six one-minute pulses, with cooling on ice in between. The cooled lysate was centrifuged at 4600xg for 5 min at 4°C, and the supernatant was then transferred to Eppendorf tubes and cleared by centrifugation at 20,000xg for 20 min at 4°C.

Purification of labelled proteins was then performed by copper-catalyzed coupling to an alkyl-agarose resin. For each sample, 200μL Click-iT® Enrichment Resin was washed once with RNase-free water and re-suspended in 1ml 2X Copper Catalyst Solution (freshly prepared according to manufacturer’s datasheet). The lysate was then added and incubated at room temperature on a rotating wheel overnight. Subsequently, resin bound proteins were reduced and alkylated as outlined by the manufacturer. For stringent removal of non-specifically bound proteins, 10 washes were performed with each of the SDS was buffer, 8 M urea and 20% acetonitrile. To release resin-bound proteins, the resin was then re-suspended in 1ml digestion buffer (100 mM Tris, 2 mM CaCl2, 10% acetonitrile), pelleted and aspirated to leave ∼200ml digestion buffer and supplemented with 10 μL of 0.1 μg/μL trypsin. This was incubated at 37°C overnight, centrifuged to pellet the resin and the digest supernatant retained.

To prepare the digest for MS, the sample was acidified to pH 1-2 using 10% TFA and processed onto activated stage-tips. Briefly, three C-18 discs (Sigma-Aldrich) were placed in a 200 μL pipet tip, and washed sequentially with 50 µl methanol, 50 µl 80% acetonitrile/0.1% TFA and 70µl 0.1% TFA. The sample was then passed by centrifugation at 1300 rcf for 5 min, washed with 70 µl 0.1% TFA and stored at –20°C.

Total protein samples were prepared simultaneously from the lysate. Approximately 25µg input protein was mixed with 1x NuPAGE LDS sample buffer supplemented with 10mM DTT, loaded onto a 4%–12% NuPAGE™ Bis-Tris Mini Protein Gel and run in MOPS buffer at 190V. Subsequently, the gel was washed with water, stained with Coomassie Protein Stain for 1 h, rinsed several times with water to destain. Protein smears were excised and resulting gel fragments were destained further by two washes in 1:1 50 mM ammonium bicarbonate (ABC) and 50% acetonitrile (ACN) for 30 min at 37°C with shaking at 1200 rpm.

Proteins were then reduced and alkylated in situ. Gel fragments were covered with 10 mM DTT in 50mM ABC for 30 min at 37°C, then shrunk by 5 minutes in ACN. Following this, fragments were incubated with 55 mM iodoacetamide in ABC for 20 min at ambient temperature in the dark, washed with ABC and shrunk twice with ACN. These were then digested overnight with 13 ng/μL trypsin (Pierce) in 10mM ABC containing 10% (v/v) ACN at 37°C.

Following trypsin digestion, samples were acidified and passed on to stage-tips as outlined for eluates. To remove any remaining peptides from the gel, the fragments were incubated for 10 min in 100% ACN, which was then transferred to a 2 mL Protein LoBind tube and dried under vacuum centrifugation at 60°C. The resulting protein pellets were resuspended in 100 μL of 0.1% TFA and passed through the stage tip. These were then washed with 100 μL 0.1% TFA, and placed at −20°C prior to mass spectrometry.

#### Mass Spectroscopy

Peptides were eluted in 40 μL of 80% acetonitrile in 0.1% TFA and concentrated down to 1 μL by vacuum centrifugation (Concentrator 5301, Eppendorf, UK). The peptide sample was then prepared for LC-MS/MS analysis by diluting it to 5 μL by 0.1% TFA.

LC-MS analyses were performed on Orbitrap Fusion™ Lumos™ Tribrid™ Mass Spectrometer (Thermo Fisher Scientific, UK) on a Data Independent Acquisition (DIA) mode, coupled on-line, to an Ultimate 3000 HPLC (Dionex, Thermo Fisher Scientific, UK). Peptides were separated on a 50 cm (2 µm particle size) EASY-Spray column (Thermo Scientific, UK), which was assembled on an EASY-Spray source (Thermo Scientific, UK) and operated constantly at 55oC. Mobile phase A consisted of 0.1% formic acid in LC-MS grade water and mobile phase B consisted of 80% acetonitrile and 0.1% formic acid. Peptides were loaded onto the column at a flow rate of 0.3 μL min-1 and eluted at a flow rate of 0.25 μL min-1 according to the following gradient: 2 to 40% mobile phase B in 150 min and then to 95% in 11 min. Mobile phase B was retained at 95% for 5 min and returned back to 2% a minute after until the end of the run (190 min).

MS1 scans were recorded at 120,000 resolution (scan range 350-1650 m/z) with an ion target of 5.0e6, and injection time of 20ms. MS2 was performed in the orbitrap at a resolution of 30,000 with a scan range of 200-2000 m/z, maximum injection time of 55ms and AGC target of 3.0E6 ions. HCD fragmentation was utilised with stepped collision energy of 25.5, 27 and 30 ^75^. Variable isolation windows were used throughout the scan range, ranging from 10.5 to 50.5 m/z. Narrower isolation windows (10.5-18.5 m/z) were applied from 400-800 m/z and then gradually increased to 50.5 m/z until the end of the scan range. The default charge state was set to 3. Data for both survey and MS/MS scans were acquired in profile mode.

The mass spectrometry proteomics data have been deposited to the ProteomeXchange Consortium via the PRIDE partner repository with the dataset identifier PXD064070.

#### RNAseq

RNA sequencing for cells growing glucose and after nutrient shift was performed as described^38^, and datasets are deposited in GEO (detailed in Table S4). BY4741 cells were grown to OD_600_ 0.45 in 50ml SC -TRP media supplemented with 2% sucrose. Sucrose grown samples were pelleted by centrifugation at 4,600 rpm for 2 minutes at 4°C, snap frozen in a dry-ice ethanol bath, and stored at -80°C. For depletion, cells were collected by filtration and transferred to SC - TRP media supplemented with 2% glycerol and 2% ethanol. Following 16 min, cells were collected and frozen as described for sucrose grown samples. Each condition was collected in biological triplicate.

To extract RNA, cell pellets were re-suspended in TRIzol reagent supplemented with 5 mM DTT final, transferred to 2 ml screw-cap tubes containing 200 μL of zirconia beads and lysed by three cycles (6m/s for 40 sec) in a Fast Prep-24 machine (MP Biomedicals). RNA was cleaned twice with chloroform and precipitated using 0.8 vol isopropanol. Samples were treated with DNase I and further purified using RNA Clean and Concentrator kit.

Libraries for RNAseq were prepared by the Wellcome Trust Clinical Research Facility at Western General Hospital (Edinburgh, UK) using the NEBNEXT Ultra II Directional RNA Library Prep kit (NEB #7760) and the Poly-A mRNA magnetic isolation module (NEB #E7490) according to the provided protocol. The libraries were sequenced using the NextSeq 2000 platform (Illumina Inc, #20038897), with paired-end, 2x 50 nt outputs.

### Quantification and Statistical Analysis

#### RNAseq data analysis

For the RNAseq experiments reported here, raw sequencing reads were first processed using Flexbar to remove Illumina adapters and low-quality bases. Reads less than 18nt or with quality threshold below 30 were discarded. These trimmed and filtered reads were aligned to the *S. cerevisiae* transcriptome using Salmon, and the reads mapping to each transcript quantified. For this, the GTF file alongside fasta files for genes, cDNAs, ncRNA were downloaded from Ensembl using genome version GCF_000146045.2_R64-1-1. These were input to the generateDecoyTranscriptome.sh Salmon script, producing a hybrid fasta containing the decoy sequences from the genome concatenated with the transcriptome. Indices for this decoy-aware transcriptome were then generated using the salmon indexer. The Salmon quant command was used to quantify the processed reads directly against this index.

Following processing, these new datasets were combined with our previous published RNAseq datasets generated by the same pipeline. The resulting data were analyzed in R. Quantified reads were normalized in counts per million (CPM) and filtered using the edgeR package, applying the TMM method and ‘filterByExp’ function (threshold per sample: CPM > 15; threshold over all samples: CPM > 20) (Table S6). Subsequently, to calculate fold changes (FC) in transcript abundance between conditions, differential expression analysis was performed in edgeR. The ‘model.matrix’ function was used to model experimental design, dispersion estimates were obtained using ‘estimateDisp’, fit to a linear model using ‘glmQLFit’ and expression analysis performed using ‘glmQLFTest’ (Table S7). The clusterProfiler package was used to analyze and visualize the overrepresentation of GO Biological Process annotations among differential expressed transcripts.

For visualization, pheatmap was used to create Spearman correlation heatmaps and cluster analysis comparing RNAseq experiments. Principle component analysis (PCA) was performed using the log transformed CPM data and plotted using biplot. Scatter graphs comparing logCPM distribution between experiments and volcano plots visualizing differentially expressed genes were created using ggplot2.

#### CRAC analysis

Our previous published CRAC datasets were re-analyzed here to address novel questions (Fig 4A). These are for the RNA interactomes of the DEAD-box helicase eIF4A and associated factor eIF4B in glucose grown and glucose starved (16 min) *S. cerevisiae.* Utilized data is outlined in Table S4. Processing, mapping and quantification of the sequencing data was described in ^16^. These reposited datasets were imported into R for analyses presented here.

Firstly, a high-confidence dataset of quantified mRNA transcripts bound under each condition was created (Table S8). Feature counts were normalized to library size in Reads per Million (RPM) and a threshold of RPM < 0.5 (∼1 count) applied. Features were required to have been quantified in a minimum of two replicates, and the median of replicate values was used. Accounting for low coverage in glucose withdrawal conditions, imputation of the threshold 0.5 RPM was used to retain features present in the averaged glucose data. Retained features were then filtered by annotation as ‘protein_coding; exon’ to select mRNA transcripts, and utilizing RNAseq data for corresponding conditions lowly expressed transcripts were removed by a RPKM threshold of 10. For each protein, changes in RNA binding before and after stress were compared using scatterplots created in ggplot2, using the log transformed RPM values. For eIF4B, RPM binding was further normalized by expression using the RNAseq RPKM values at the equivalent timepoint. As mRNA populations will not alter significantly in seconds, RNAseq for glucose was used for 30 sec withdrawal. Targeted analyses of changes in eIF4B binding were performed for the top 100 most increased (induced) and decreased (repressed) transcripts following glucose withdrawal, as determined by differential expression analyses of RNAseq data (outlined above; Table S9). Bar graphs were created using GraphPad Prism 10.

In order to visualize binding across individual transcripts, the coverage at each position along the genome was calculated and normalized to the library size using genomecov from bedtools v2.27.0 ^76^. The Integrative Genomics Viewer was used to illustrate the distribution of reads across chosen transcripts ^77^.

#### Metabolic Labelling Proteomics Analysis

The DIA-NN software platform version 1.9.2. was used to process the DIA raw files, and these were searched against the *Saccharomyces cerevisiae* complete/reference proteome in Uniprot (released in June, 2019). Precursor ion generation was based on the chosen protein database (automatically generated spectral library) with deep-learning based spectra, retention time and IMs prediction. Digestion mode was set to specific with trypsin allowing maximum of two missed cleavages. Carbamidomethylation of cysteine was set as fixed modification. Oxidation of methionine, and acetylation of the N-terminus were set as variable modifications. The parameters for peptide length range, precursor charge range, precursor m/z range and fragment ion m/z range as well as other software parameters were used with their default values. The precursor FDR was set to 1%. Annotating library proteins were created with information from the FASTA database: the spectral library contained 6093 proteins and 6093 genes.

Processed protein intensity data was analyzed in R. Standard filtering steps were applied to clean the dataset, and reverse and potential contaminant were removed. The dataset contained triplicate samples of the labelled input (total) and eluate (nascent) proteins for AHA-purification (described above), alongside unlabeled controls. To reliably identify the nascent proteome, only protein groups quantified in all purification eluate samples for at least one labelled condition were retained (n=4165).

Samples were VSN normalized and missing values were imputed with the left-censored MinProb method, as proteins are not expected to be missing at random, using the proteomics specific DEP package. Subsequently, to quantify changes between conditions, differential expression tests were performed using test_diff function on manually defined contrasts. Obtained *p* values were corrected to control for false discoveries using the highly stringent Bayesian/local false discoveries rate (FDR) method, and proteins with greater than 1.5-fold change (FDRL≤L0.05) were considered differentially expressed. Volcano plots visualizing differentially expressed genes were created using ggplot2.

For cluster analysis of nascent protein production, proteins with significant change between +AHA purification eluates were selected using the conventional FDR in R stats. For the resulting 524 proteins, median intensities for the input and eluate samples were z-scored and k-means clustered (k=4). For visualization, pheatmap was used to create correlation heatmaps. Clusters 2 and 3 showed significantly increased label incorporation in glucose or no glucose conditions, respectively, and were analyzed further. For genes in these clusters, RNAseq and CRAC data sets were utilized to compare fold change in RNA abundance (RPKM), eIF4B binding (RPM) and protein abundance (intensity) following glucose withdrawal.

## SUPPLEMENTARY FIGURE LEGENDS

**Supplementary Figure 1.**
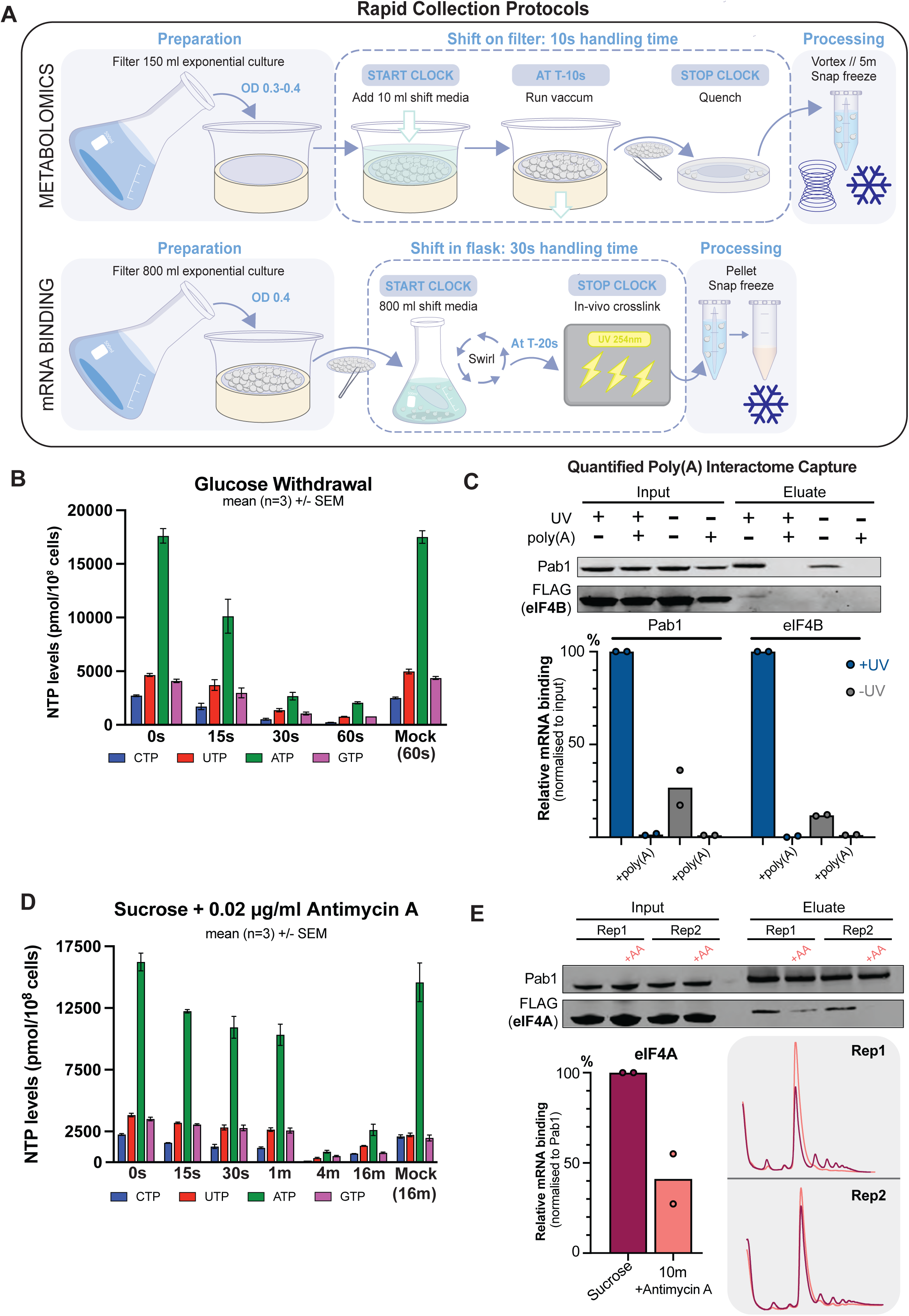
Sample preparation and effects of glucose withdrawal (related to Figure 1) A: Schematic overview of the processes used for rapid carbon-source shift experiments. For metabolomics, exponential cultures were harvested by vacuum filtration. The fresh medium was applied while the yeast was retained on the filter and incubated for the indicated time. 10 sec was required to remove media and transfer the filter to lysis solution. In poly(A) interactome capture experiments, filtered cells were transferred to a fresh flask of shift media, resuspended and incubated shaking. The shifted culture was UV crosslinked at 254nm to stabilize RNA-protein interactions. 20 sec was required for handing and crosslinking. Yeast was then re-harvested for processing. Steps following crosslinking are less time-critical, as the covalent bonds have already been generated. B: Bar chart showing NTP levels measured following glucose withdrawal in *S.cerevisiae.* Replicate samples (n=3) were obtained at 15 sec, 30 sec and 1 min following shift from 2% glucose to 2% glycerol/ethanol or a mock shift back to glucose. Error bars represent the mean +/− standard error (SEM). C: Assessment of poly(A)-interactome capture specificity using non-crosslinked and poly(A) competed controls. Exponential glucose grown cultures were either UV crosslinked at 254 nm for 12s (+UV) or left untreated (-UV) prior to denaturing cell lysis. Diluted lysates (15mg/ml) were either input directly to pulldowns or prepared with artificial poly(A) (0.4mg/ml) as a competitor. The amount of Pab1and FLAG-tagged eIF4B purified and eluted in each case was assessed by western blotting (upper) and quantified by intensity relative to the input level (n = 2). Bar graphs (lower) represent the amount of Pab1 or eIF4B obtained in the eluate (mRNA-bound) as a percentage of the input level. UV treated samples are shown in blue, compared to untreated samples in grey. D: Bar chart showing NTP levels measured following Antimycin A (AA) treatment in yeast grown on sucrose. Replicate samples (n=3) were obtained at 15 sec, 30 sec, 1 min, 4 min and 16 min following shift to 2% sucrose media containing 0.02µg/ml AA or a mock return to 2% sucrose. Error bars represent the mean +/-standard error (SEM). E: Simultaneous assessment of eIF4A-mRNA binding and translational status following AA treatment. To allow polysome profiling and poly(A) interactome capture of the same culture, prior to or following shift from 2% sucrose to equivalent media containing 0.02µg/ml AA (10 min), these were split. The result of two distinct biological experiments is shown. Association of FLAG-tagged eIF4A with mRNA was assessed by poly(A)-interactome capture and western blotting (top), and quantified by relative intensity normalized to poly(A) binding protein (Pab1) as a control for input and pull-down efficiency. Bar graph shows the amount of eIF4A bound to mRNA as a percentage of the pre-AA treatment level, the corresponding polysome gradient analyses are shown (grey box) with pre-treatment in scarlet and +10m AA in peach.

**Supplementary Figure 2.**
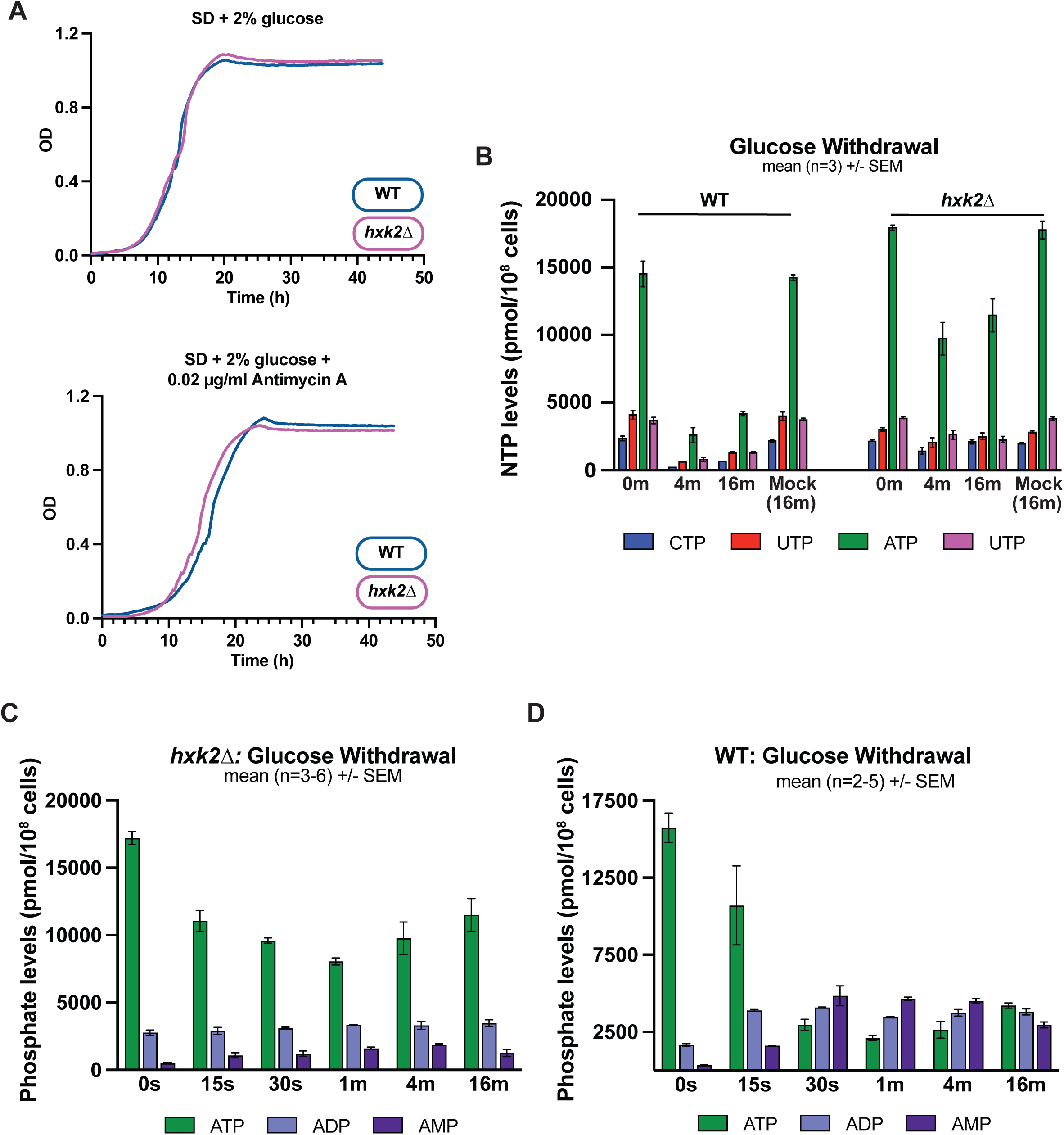
Nucleotide levels in wildtype and strains lacking Hxk2 (related to Figure 2) A: Growth curves comparing growth of wild-type (WT, blue) and *hxk2*Δ (pink) BY4741 yeast strains. Growth rate on 2% glucose (SD) alone (upper) and following addition of 0.02µg/ml Antimycin A to exponential cultures (lower) was assessed by change in optical density (OD) over 48 hours. n=2. B: Bar chart showing NTP levels measured following glucose withdrawal over minute time-courses in wild type (WT) and *hxk2*Δ strains. Replicate samples (n=3) were obtained at 0 min, 4 min, and 16 min following shift from 2% glucose to 2% glycerol/ethanol or a 16 min mock shift back to glucose. Error bars represent the mean +/− standard error (SEM). C: Bar chart showing levels of intracellular adenosine phosphates in *hxk2*Δ yeast (monophosphate, AMP; diphosphate, ADP; triphosphate, ATP). Samples were obtained at 0 min, 15 sec, 30 sec, 1 min, 4 min and 16 min following shift from 2% glucose to 2% glycerol/ethanol or a 16 min mock shift back to glucose. Each condition has 6 replicates, except for 15 sec where n=3. D: As in C for WT. Each condition has 5 replicates, expect for 15 sec where n=2. Error bars represent the mean +/− standard error (SEM).

**Supplementary Figure 3.**
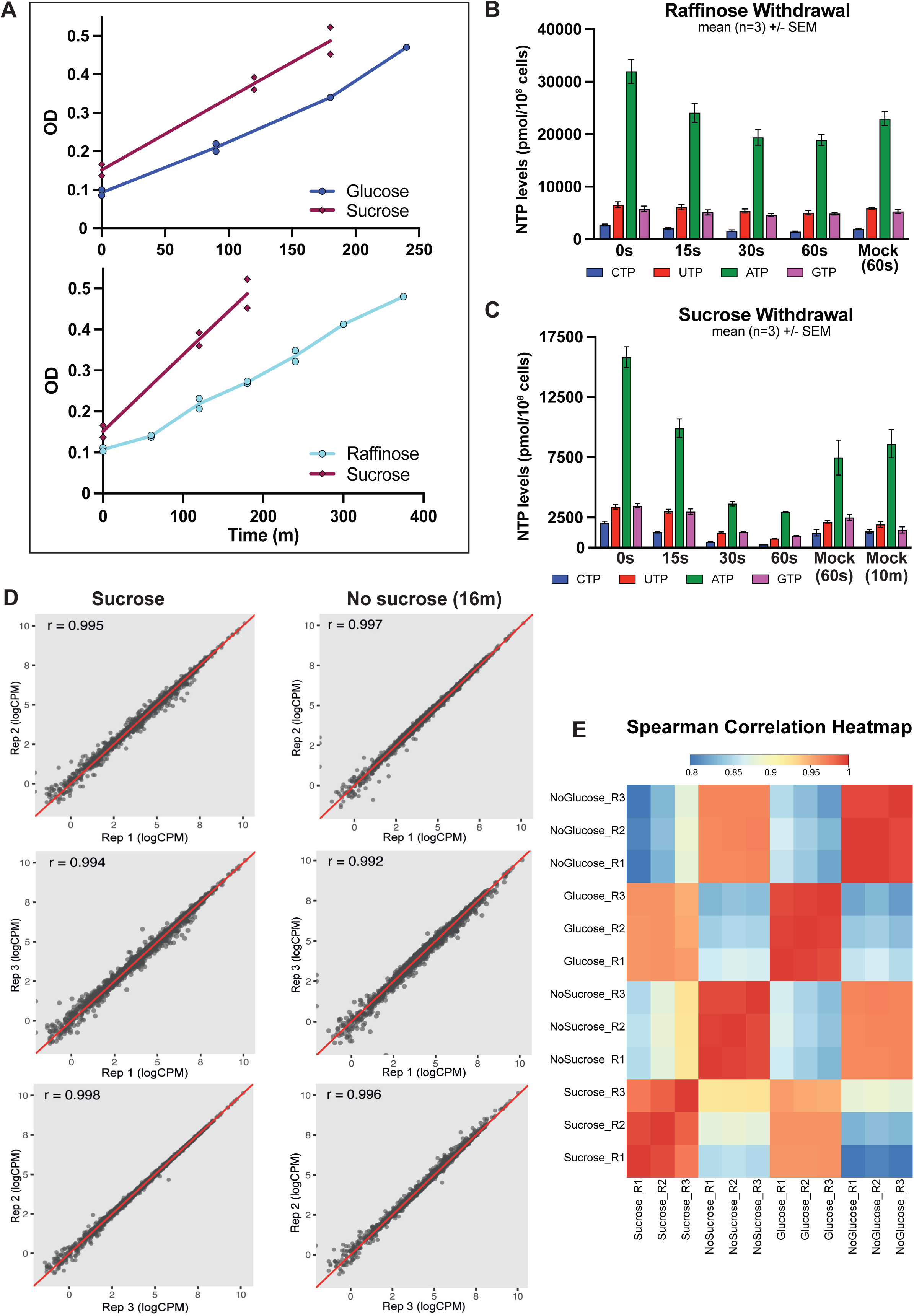
Comparison of withdrawal of different carbon sources (related to Figure 3) A: Comparison of growth rate in yeast utilizing 2% sucrose relative to 2% glucose (upper) or 2% raffinose (lower). Growth was assessed by change in optical density (OD) during the exponential phase (from OD_600_ 0.1 - 0.5). n=2. B: Bar chart showing NTP levels measured following raffinose withdrawal in *S.cerevisiae*. Replicate samples (n=3) were obtained at 15 sec, 30 sec and 1 min following shift from 2% raffinose to 2% glycerol/ethanol or a mock shift back to raffinose media. Error bars represent the mean +/− standard error (SEM). Mock shifts resulted in mild ATP depletion, but levels remain well above the Km for eIF4A after 60 sec C: Bar chart showing NTP levels measured following sucrose withdrawal in *S.cerevisiae*. Replicate samples (n=3) were obtained at 15 sec, 30 sec and 1 min following shift from 2% sucrose to 2% glycerol/ethanol. Mock shift samples were harvested equivalently and returned to sucrose media for indicted times. Error bars represent the mean +/− standard error (SEM). D: Scatter plots comparing replicate RNAseq datasets for sucrose grown and sucrose starved (16 min) *S.cerevisiae.* Counts are normalized for library depth as counts per million (CPM), all mRNAs for which > 15 sequence counts were recovered across 3 replicates are displayed. The correlation coefficient (r) for each replicate pair is shown. E: Heatmap showing spearman correlation of sucrose withdrawal RNAseq datasets, with equivalent available data for glucose withdrawal ^38^. Samples were obtained prior-to or following shift from media containing either 2% glucose or sucrose to 2% glycerol/ethanol for 16 min (NoGlucose and NoSucrose, respectively). Spearman correlations between RNAseq data replicates were calculated using all mRNAs for which > 15 sequence reads were recovered (n=5,028).

**Supplementary Figure 4.**
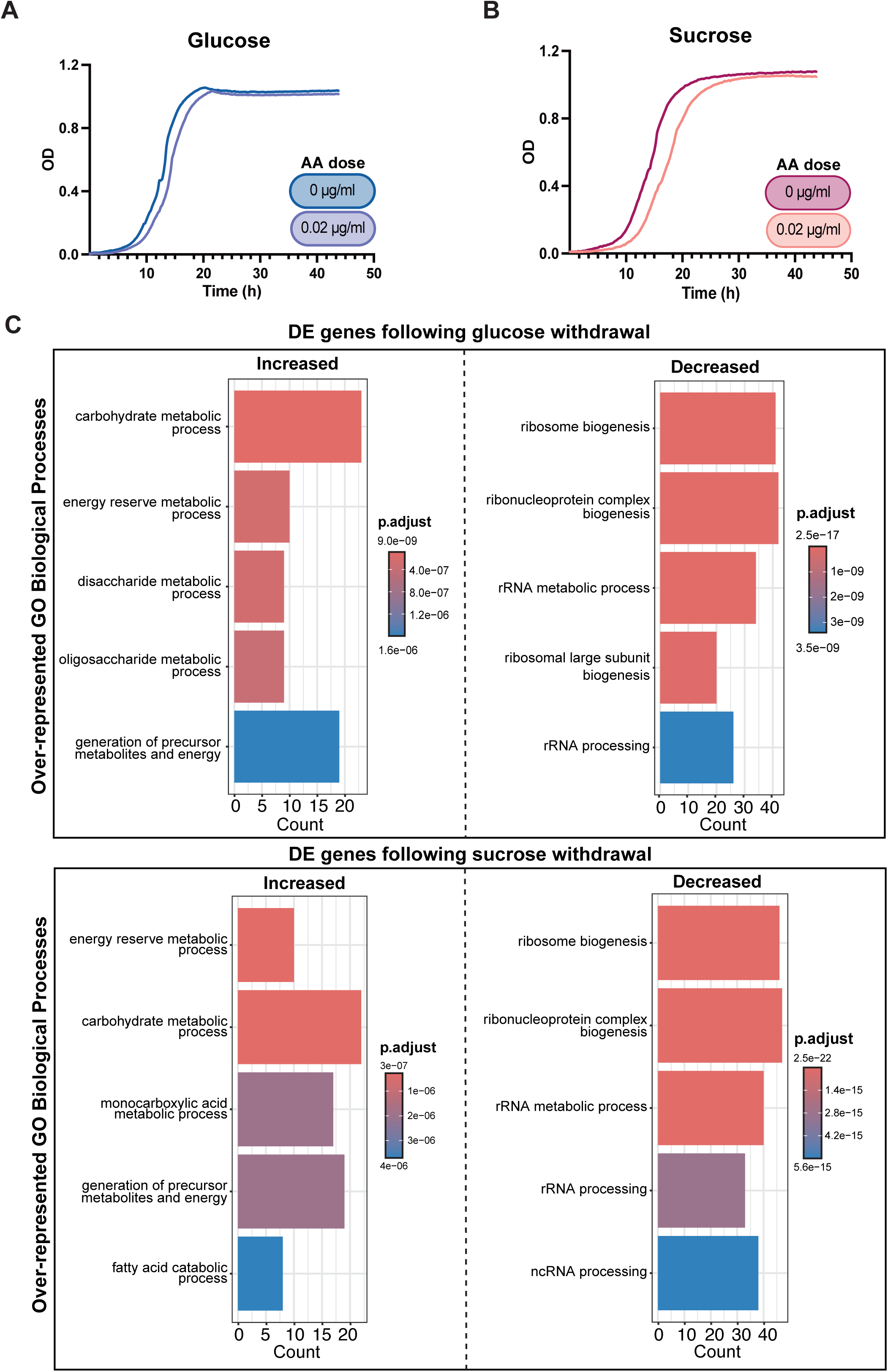
Transcription following glucose or sucrose withdrawal (related to Figure 4) A: Growth curves comparing sensitivity to the mitochondrial inhibitor Antimycin A (AA) in yeast grown on glucose. The effect of the addition of 0.02µg/ml AA to the growth rate of exponential cultures was assessed by change in optical density (OD) over 48 hours as compared to mock treated cells (0µg/ml). n=2. B: As in A, for yeast utilizing sucrose as carbon source. C: Functional analysis of transcriptional changes following withdrawal of glucose (top) or sucrose (bottom) as carbon source. Gene ontology (GO) over-representation analysis was performed on the 100 transcripts with the greatest fold-change in expression (FDR <0.05) following withdrawal in either case. The 5 most enriched biological processes are shown, ordered and colored by significance of enrichment, given by adjusted p value (p.adjust). The number of transcripts annotated to each term is displayed (Count).

**Supplementary Figure 5.**
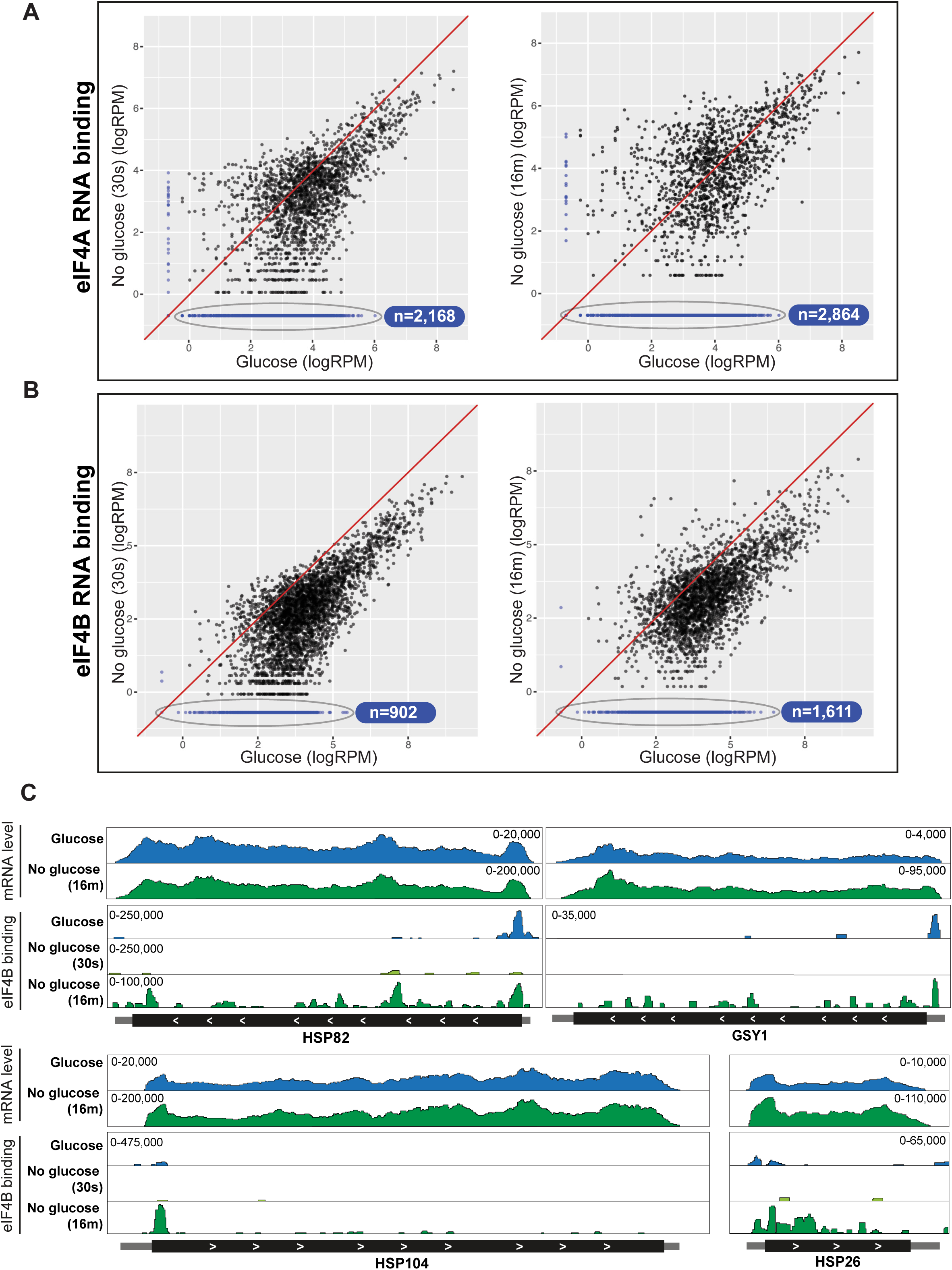
Binding of eIF4A and eIF4B after stress (related to Figure 5) A: Scatter plots comparing mRNA binding by eIF4A following either 30 sec (left) or 16 min (right) glucose withdrawal to glucose replete conditions. Transcript counts are normalized to library size, in Reads per Million (RPM), and filtered for RPM > 0.5 in all the replicates of at least one condition (n =4,377). For transcripts detected in only one condition, imputation of the minimum threshold (RPM =0.5) was used, these are colored blue and ringed, with population size indicated. B: As in A, for eIF4B C: Examples of eIF4B binding across transcripts induced following glucose withdrawal (HSP82, GSY1, HSP104, HSP26), as in 5F.

**Supplementary Figure 6.**
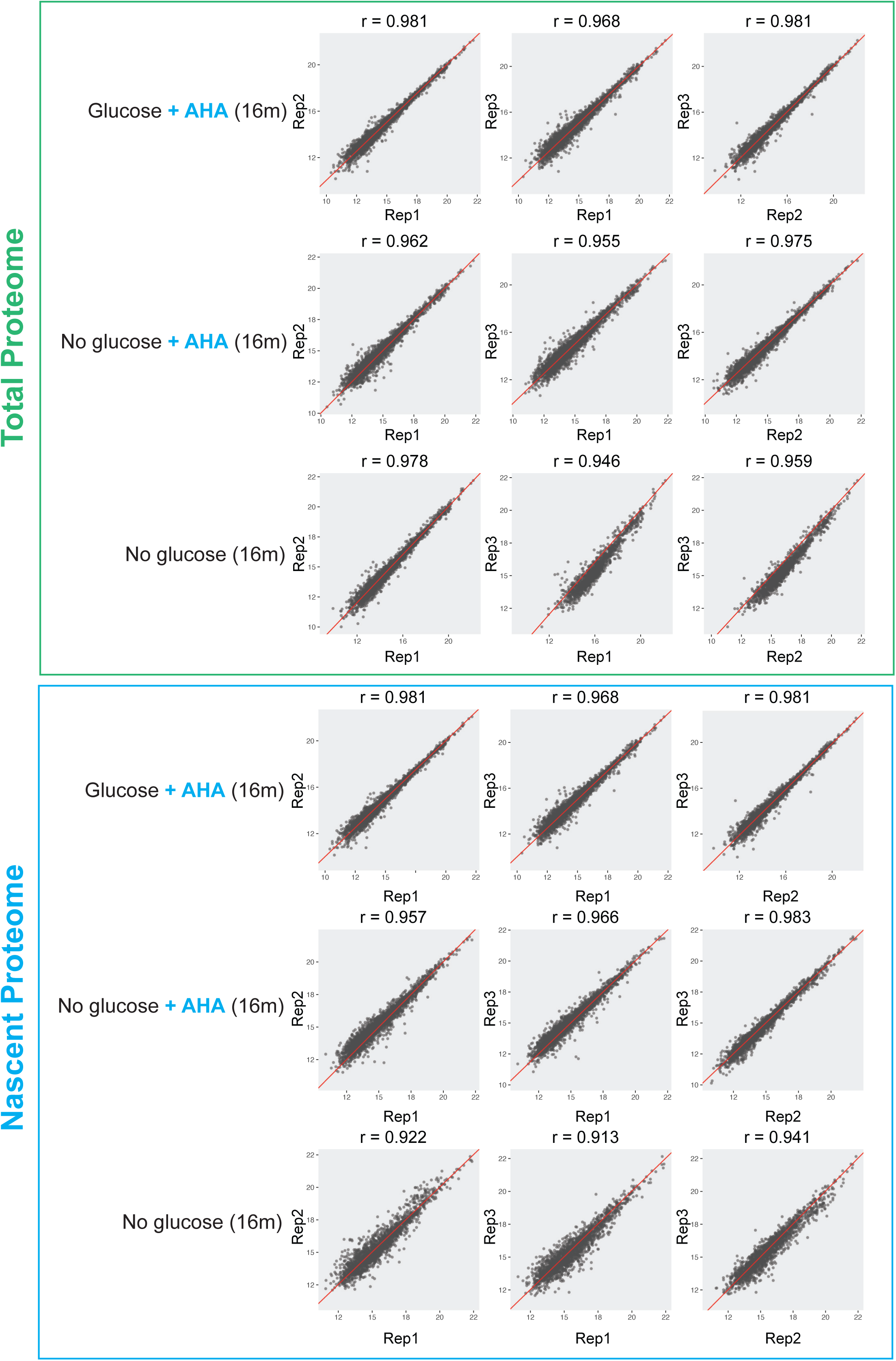
Comparison of proteome labeling replicates. (related to Figure 6) Scatter plots comparing replicate proteomic data sets. Yeast cultures were labelled using the methionine analogue L-Azidohomoalanine (AHA) for 16m, either directly in 2% glucose media or alongside shift to 2% glycerol/ethanol (No glucose). Unlabeled controls were prepared by transfer to media lacking AHA for the equivalent time. Nascent proteomes (blue box) were obtained by purification of AHA-labelled proteins see Figure 6a), while total proteomes reflect the inputs (green box). Protein intensities are log2 normalized, and the correlation coefficient (r) for each replicate pair is shown.

**Supplementary Figure 7.**
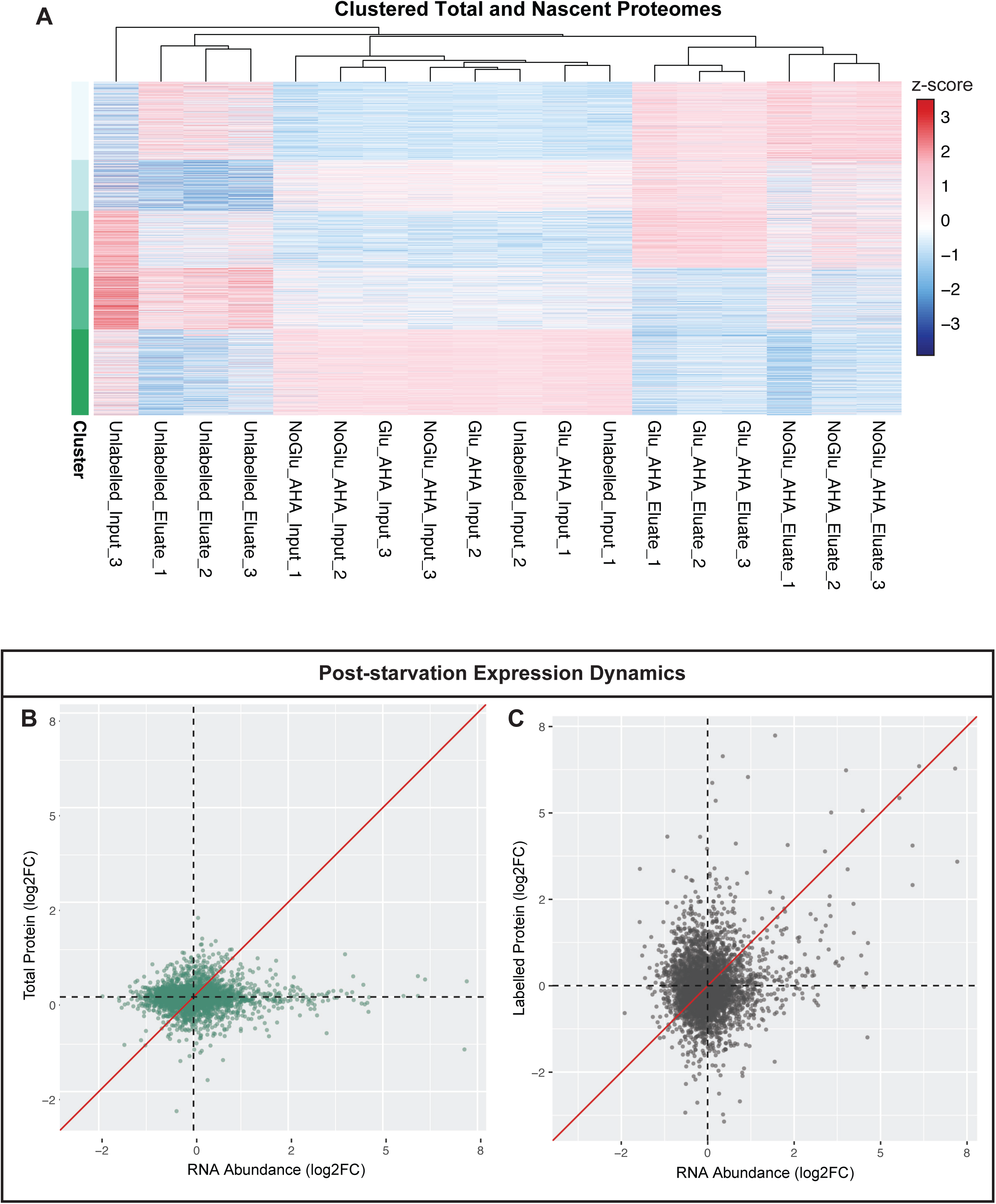
Clustering of nascent proteomes. (related to Figure 7) A: Heatmap for all proteomes analyzed (as described S6), comparing quantified protein intensity by z-score (deviation from average). K-means clustering was used to define sets of proteins with similar expression profiles (k = 5, 100 rounds) (rows). Within these groups, samples were ordered by hierarchical clustering, visualized by accompanying dendrogram (columns). B: Scatter plots comparing fold change (FC) in abundance (RPKM) of a transcript following 16 min glucose withdrawal, to the level of the associated protein in the nascent (labelled, left) and total (right) proteome. Fold change (FC) in log2 normalized protein intensities are plotted, and the x=y line is shown in red.

## DATA TABLES

S1: Deposited Data Sets

S2: Quantified NTP Levels Reported as Estimated Concentrations

Average of 3 replicates in pmol/cell, then converted to intracellular concentrations (mM) as described in methods.

S3: Global Changes in RNA Levels during Withdrawal.

WT Glucose, WT No Glucose (16 min), WT Sucrose and WT No Sucrose (16 min), datasets were combined following equivalent filtering, trimming and alignment. Quantified reads were normalized in CPM and filtered using edgeR TMM method (threshold per sample: CPM > 15; threshold over all samples: CPM > 20).

S4: RNAseq Differential Expression Analysis Results

Showing Fold Change between conditions for all transcripts, on a log scale. Significance is given by false discovery rate.

S5: CRAC Results Showing the Amount of Binding (in Reads per Million) to Different mRNAs for eIF4B and eIF4A

Quantified mRNA transcripts bound under each condition. RPM normalized counts, threshold of RPM < 0.5 (∼1 count) applied. Features were required to have been quantified in a minimum of two replicates and the median of replicate values was used. Imputation of the threshold 0.5 RPM was used to retain features present in the averaged glucose data. Filtered by annotation as ‘protein_coding; exon’ to select mRNA transcripts. RNAseq data - RPKM threshold of 10.

S6: Top Induced and Repressed Transcripts Following Sugar Withdrawa**l**

The 100 most increased and decreased transcripts (FDR < 0.05), respectively, following 16 min glucose withdrawal. These are ranked by magnitude of fold change. eIF4B mRNA binding (RPM) quantified by CRAC for each transcript is provided.

S7: L-Azidohomoalanine (AHA) Metabolic Labelling Proteomics Data

Normalized protein intensities (log2) for the input and eluate of purification of AHA-labelled proteins. These reflect the total and nascent proteome, respectively. Proteomes were VSN normalized and imputed by the ‘MinProb’ method.

S8: Cluster analysis of nascent protein production, related to Figure 7.

Identity of clusters allocated by K means clustering (k=4) of z-scored median intensities for 524 proteins showing differential expression between AHA purification eluates.

S9: Proteomics Differential Expression Analysis Results, related to Figure 6 and 7. Outcome of analysis using DEP for indicated comparisons. Proteins identified as differentially expressed (FC > 1.5, FDR <0.05) are included.

